# A 21-year survey of *Escherichia coli* from bloodstream infections (BSIs) in a tertiary hospital reveals how community-hospital dynamics of the B2 phylogroup clones influence local BSI rates

**DOI:** 10.1101/2020.04.10.034777

**Authors:** Irene Rodríguez, Ana Sofia Figueiredo, Melissa Sousa, Sonia Aracil-Gisbert, Miguel Díez Fernández de Bobadilla, Val F. Lanza, Concepción Rodríguez, Javier Zamora, Elena Loza, Patricia Mingo, Claire J. Brooks, Rafael Cantón, Fernando Baquero, Teresa M Coque

## Abstract

This is a longitudinal study comprising 649 *Escherichia coli* (EC) isolates representing all 7165 EC-BSI episodes recorded in a hospital (1996-2016). Strains analysis included clonal identification (phylogenetic groups/subgroups, STc131 subclades, PFGE, and WGS), antibiotic susceptibility (13 antibiotics), and virulence-associated genes (VAGs, 29 genes). The incidence of BSI-EC increased from 1996 to 2016 (5.5 to 10.8 BSI episodes/1000 hospitalizations, average 7-8/1000). B2 isolates predominate (53%), subgroups B2-I (STc131), B2-II, B2-IX, and B2-VI representing 25%, 25%, 14%, and 9%, respectively. Intertwined waves of community-acquired (CA) + healthcare-associated and community-onset healthcare-associated (HCA), and hospital-acquired (HA) episodes of both B2 and non-B2 phylogroups occurred. A remarkable increase was only observed for B2-I-STc131 (C1/C2 subclades), with oscillations for other B2 subgroups and phylogroups throughout the years. Epidemic and persistent clones (comprising isolates with highly similar/identical-PFGE types and genomes differing in 18-97 SNPs) of B2-I (STc131), B2-II (STc73), B2-III (STc127), B2-IX (STc95), and B2-VI (STc12) were recovered from different patients, most at hospital admission, for long periods (2-17 years), ESBL producers or resistance to ciprofloxacin in B2 isolates were almost restricted to B2-I (STc131) subclade C. STc131 contributed to increasing the B2 rates but only transiently altered the EC-population structure.

The increase of EC-BSI was determined by waves of CA+HCA-BSI episodes that predate the waves of HA-BSI. Besides the risk of hospital transmission that led to temporal increases in BSIs, this study suggests that EC-populations/clones from community-based healthy individuals may occasionally have an epidemic structure and provide a source of transmissible strains influencing the HA-BSIs incidence.

**IMPORTANCE:** Sepsis is the third cause of mortality in Western countries and one of the Global Health threads recognized by the WHO since 2017. Despite *Escherichia coli* constitutes the most common cause of bloodstream infections (BSI), its epidemiology is not fully understood, in part due to the scarcity of local and longitudinal studies. Our work analyzes the long-term dynamics of *E. coli* causing bacteremia in a single institution and reveals waves of different clonal lineages that emerge periodically and successfully spread afterward in both the community and hospitals. Because the origin of BSI-*E. coli* infections is the gut, the microbiota of healthy individuals might occasionally have an epidemic structure, providing a source of *E. coli* strains to influence the incidence of hospital BSIs. The study complements previous fractionated observations focusing on specific *E. coli* lineages or antibiotic-resistant isolates in the last decades and helps to understand the epidemiology of *E. coli* BSIs and the dynamics of pandemic clones.

## INTRODUCTION

The increasing and progressive annual rate of BSIs episodes at a global level (9-13% in Western countries), made the WHO to consider sepsis as a Global Health Threat in 2017 (https://www.global-sepsis-alliance.org/news/2017/5/26/wha-adopts-resolution-on-sepsis). The problem affects more than 30M of people in the world and represents the third cause of mortality in Europe and North America (1–4). *Escherichia coli*, a commensal opportunistic pathogen of humans and animals, constitutes the primary cause of bloodstream infections (BSIs) (5, 6). The gut microbiota is often the origin of all extraintestinal infections caused by *E. coli* (7) and vary between humans of different age and lifestyle (7, 8). Among the 7 major phylogenetic groups of the species (A, B1, B2, C, D, E, F), the B2 is nowadays predominant in clinical and fecal isolates from adults and children of Western areas (7–9). Despite the apparent persistent structure of *E. coli* in the gut of these individuals, clonal expansions of emerging STcs periodically occurred (10, 11). Currently, several *E. coli* phylogenetic groups and some of the 10 B2-*E. coli* subgroups know (B2_I_-B2_X_), are overrepresented by certain sequence type complexes (STcs) [e.g.B2_I_ (STc131), B2_II_ (STc73), B2_IX_ (STc95), D (STc69) or F (STc648)], although their abundance and diversity vary between human populations (10, 12–14).

*E. coli* bacteremia was not considered common at the beginning of the 20th century but it has steadily increased for decades according to long surveys performed at Boston City Hospital between 1935 and 1972 (15), at St. Thomas hospital in London between 1969 and 1986 (16), and more recently, at hospitals in Western countries (most are cross-sectional or longitudinal multicentric studies) (1, 4, 17–19). The survey performed during the 1980s reflected, for the very first time, an incidence peak of BSI caused by *E. coli* coincidental with a community-based clonal outbreak caused by an *E. coli* O15:H12 which led to its introduction in the hospital setting and a subsequent increase in nosocomial BSI cases(16). Shortly afterwards, community-based and hospital clonal outbreaks by *E. coli* of phylogroup D were extensively reported during the 1990s, namely ST69 in the US and ST393 O15:H12 in the UK and Spain (20). The most comprehensive recent analysis, using European Antimicrobial Resistance Surveillance System (EARSS) data from 2002 to 2008, also highlighted a remarkable average annual increase of 29.9% in the number of reported bacteremia caused by *E. coli* isolates resistant to 3GCs (1) although phylogenomic data were not provided in that publication. Nonetheless, many studies performed during the 2000s and afterwards documented the increasing of CA outbreaks of B2 *E. coli* lineages as STc73 (B2-II), STc95 (B2-IX) and more recently, STc131 (B2-1). Currently, all them are considered “pandemic clones” due to their global predominance (13, 17). The current knowledge suggests differences between countries but such information is highly fractionated in multicentric studies (1), mostly focused on antibiotic resistant BSI isolates (21), and performed at variable periods of time using different sampling criteria (1–4, 19, 21). Only studies from the UK, where surveillance of BSI is compulsory since 2011, provide a long-term comprehensive analysis of BSI and the population structure and the dynamics of *E. coli* causing BSI (4, 17).

Unfortunately, clonal expansions at local level have been poorly analyzed and mostly under the perspective of antimicrobial resistance and for very limited periods of time. However, local settings are important sources of information because reflect the dimensions of human populations structure (age, sex, interconnectedness), and offer stability in terms of the intensity of selection (e.g. common policies to control antimicrobial resistance such as antibiotic use, infection control strategies), models of healthcare delivery, and measuring approaches (diagnostic tools), (1, 19), all these issues being important to meet the epidemiology of infectious agents (22).

We retrospectively studied a randomized sample of 649 isolates drawn from a collection of 7165 *E. coli* isolates, which represented all BSI episodes registered at our institution between 1996 and 2016. This period coincided with the global emergence and amplification of various *bla*_ESBLs_ genes and B2 *E. coli* clones and with the overall increase in the frequency of *E. coli* BSIs. The aim of this study was to infer the local diversity and dynamics of *E. coli* causing BSIs, focusing on the B2 phylogroup.

## RESULTS

### Epidemiology of the 7165 *E. coli* isolates causing BSIs at Hospital Ramón y Cajal

The incidence of BSIs caused by *E. coli* from 1996 to 2016 in our institution ranges from 5.5 to 10.8 BSI episodes/1000 hospitalizations, with fluctuations of 7-8 BSI episodes/1000 admissions most of the years studied. The highest incidence peak observed in 2016 parallels the blooming of clinical isolates producing CTXM-27 in our and other hospitals (23). Although the overall number of BSI episodes in the hospital and community settings was similar at the beginning of the study in the mid 1990s, we observed a steady increase in both hospital-acquired (HA)-BSIs and community acquired (CA)+healthcare associated (HCA)-BSIs from 1995 to 2002 (CA+HCA/HA ratio >1–2) followed by waves of alternative predominance of either CA+HCA - BSIs or HA-BSIs. The increases of BSI episodes in the community seems to predate those in the hospital setting and would explain the wave dynamics between the hospital and the community-based populations suggested in the literature and the overall shift in the ratio of BSI acquired in the community and the hospital (Figure 1). The analysis of antimicrobial resistance records in our department for the blood *E. coli* isolates revealed a coincidental increase in the rise of BSI infections and the increasing trends of *E. coli* resistant to fluoroquinolones (FQ^R^), mainly ciprofloxacin (Cip^R^), from 1994, and resistant to third-generation cephalosporins (3GC) from 2003. Rates of resistance to other antibiotics remained stable during the period of study (data not shown).

**Figure 1.**
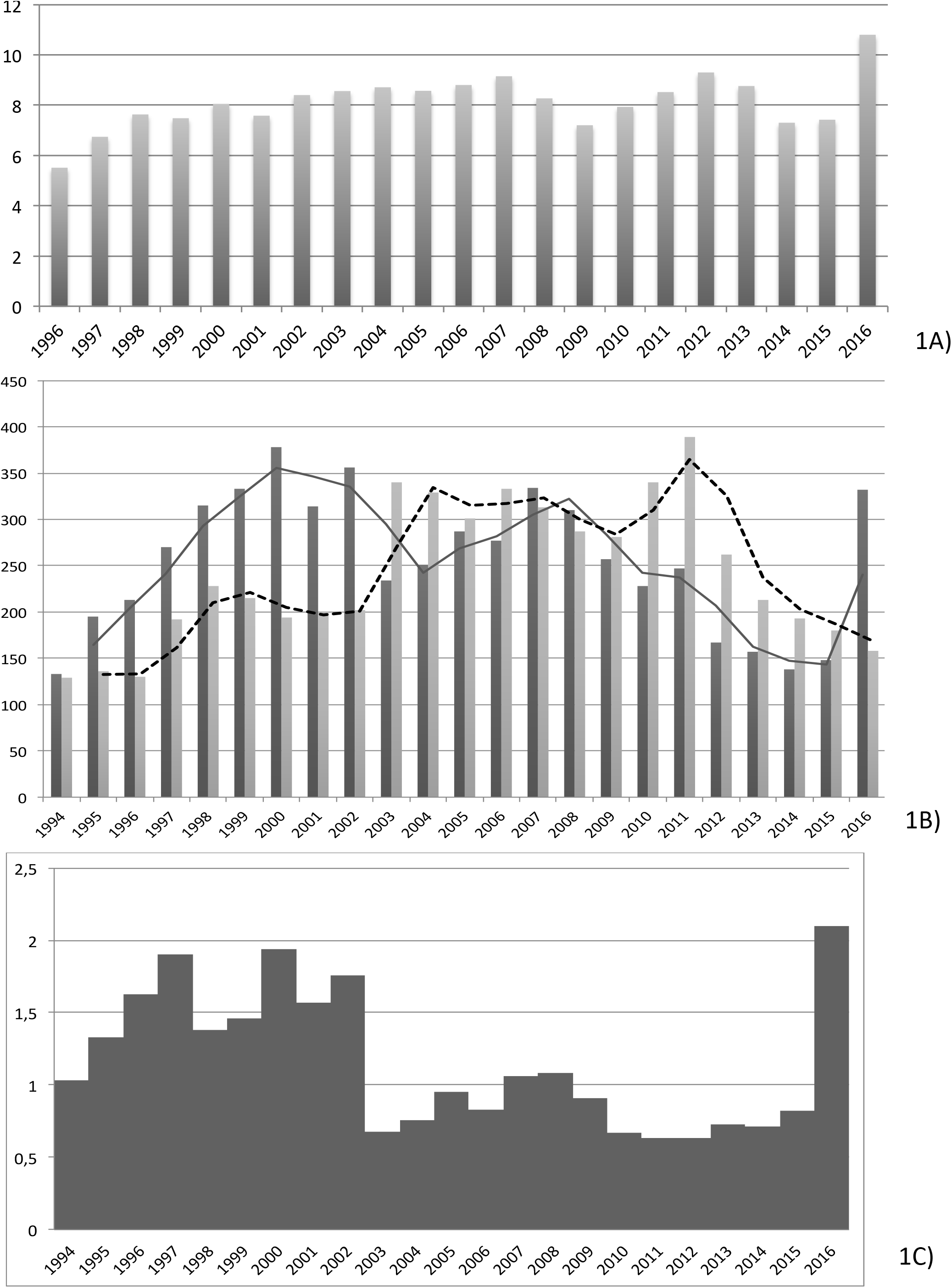
Incidence and Origin of BSI episodes caused by *Escherichia coli a*t the HRyC (1994-2016). (A) Incidence of *E. coli* BSI (episodes/1000 hospitalizations); (B) Occurrence of HCA+CA and HA episodes of BSI; (C) Ratio of HCA+CA/HA BSI episodes. Abbreviations: HA = hospital acquired; CA= community acquired; HCA = community-onset healthcare-associated. Bars in black and bars in grey represent HA episodes and CA episodes, respectively. Full and dashed lines represent the dynamics of HA episodes and CA episodes, respectively.

UTIs were identified as the origin of the BSI in one-third of the cases. This finding was more frequent in women than in men (42.33% vs. 26.77%, p≤.005) but not significantly different between patients of different age groups (p >.05). The proportion of polymicrobial BSIs was 9% (n = 57/649), and was similar for men and women (11.8% vs. 7.3% of the BSI cases, respectively).

### Clonal diversity of *E. coli* causing BSI

The predominant phylogenetic group was B2 (348/649; 53.06%), followed at a much lower frequency by D (11,4%), B1 (7,24%), A (6.3%), C (5.,1%), F (6.8%), E (0.9%) and Clades I and II (0.9%). Half of the B2 isolates corresponded to subgroups B2-I (25.6%) and B2-II (25.1%), which were followed by B2-IX (14.2%), B2-VI (9.5%) and others (Figure S1). The STc131 isolates, clearly predominant within the B2-I subgroup, represents 12.3% of the total BSI *E. coli* isolates and 21.8% of the B2 phylogroup. The STc131 isolates (serogroups O25b and O16 representing 92% and 8%, respectively) split in clade A (7/82, 9%), clade B (16/82, 20%), and clade C (59/82, 71%). Clade C comprised isolates of subclades C1 (29/82, 35.4%), C2 (22/82, 26.8%), C0 (6/82, 7.3%), and C1-M27 (2/82, 2.4%).

### Epidemiological characteristics of B2-*E. coli* isolates

#### Acquisition of the BSIs

Most B2 isolates were recovered from community-based patients although overlapping waves of CA+HCA-BSI and HA-BSI episodes of strains of both B2 and non-B2 phylogroups, which occurred until 2008, when B2 apparently overtook non-B2 BSI episodes and STc131 become transiently predominant (Figure 2 and 3). Oscillations were observed for both CA+HCA and HA episodes although an increasing trend was only detected for the HA ones (Figure 2). The stratification of the data (CA vs HCA+HA, B2 vs non-B2,..) made the sample size of each subgroup too small to infer significance. Nonetheless, the data is in agreement with the numbers obtained for all BSI cases (Figure 1).

**Figure 2.**
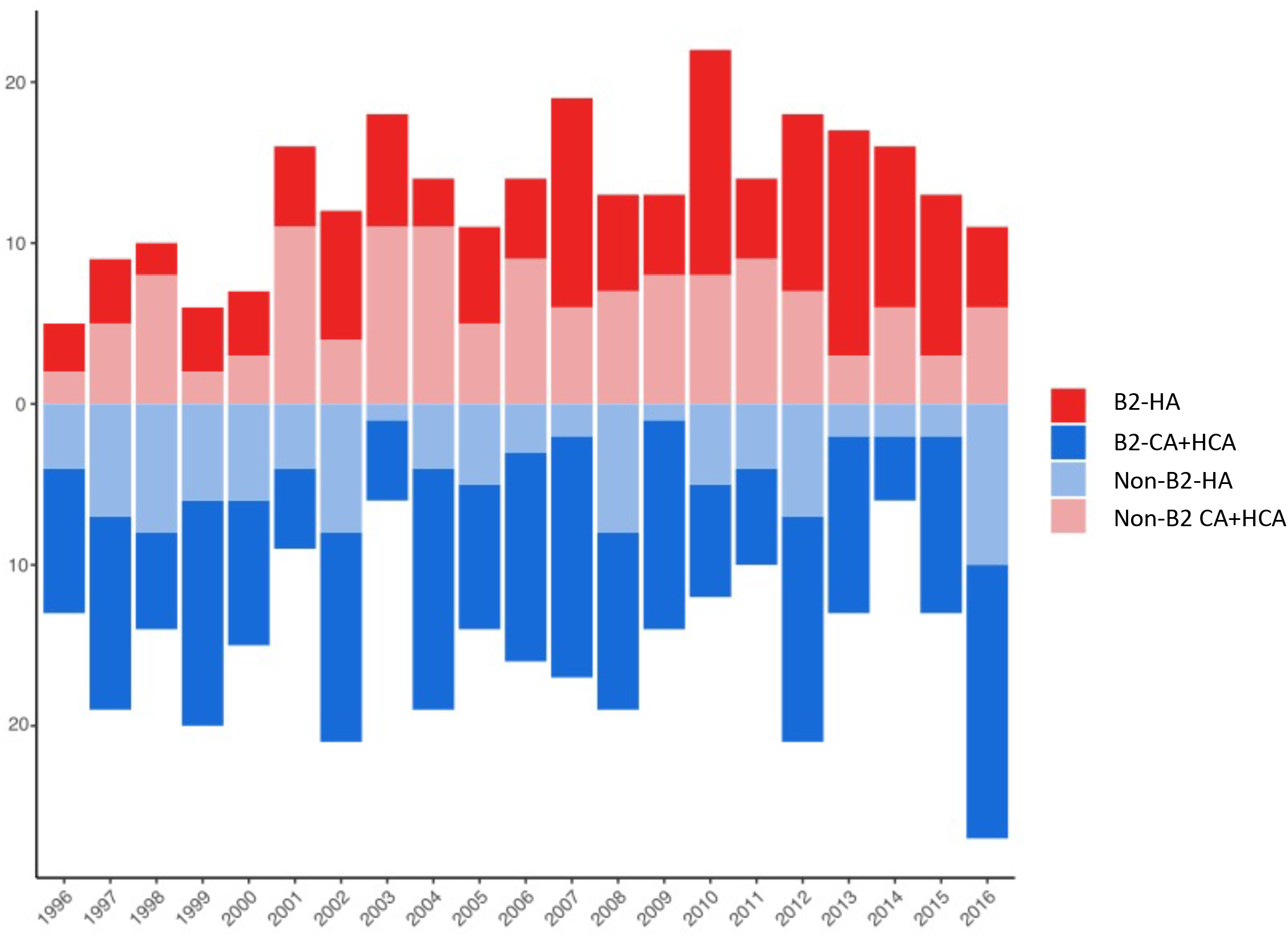
Dynamics of B2 and non-B2 lineages in hospital acquired (HA) vs community acquired (CA) + healthcare associated and community-onset healthcare-associated (CA+HCA) BSI *E. coli* isolates. Panel 2A) Stack bar plot representing the rates of B2 and nonB2 HA (bars red and light red, respectively) and B2 and non-B2 CA+HCA (bars blue and light blue, respectively). Panel 2B) Trends of the HA and CA+HCA BSI episodes represented in panel 3A. Lines red and blue correspond to HA and CA+HCA cases, respectively.

**Figure 3.**
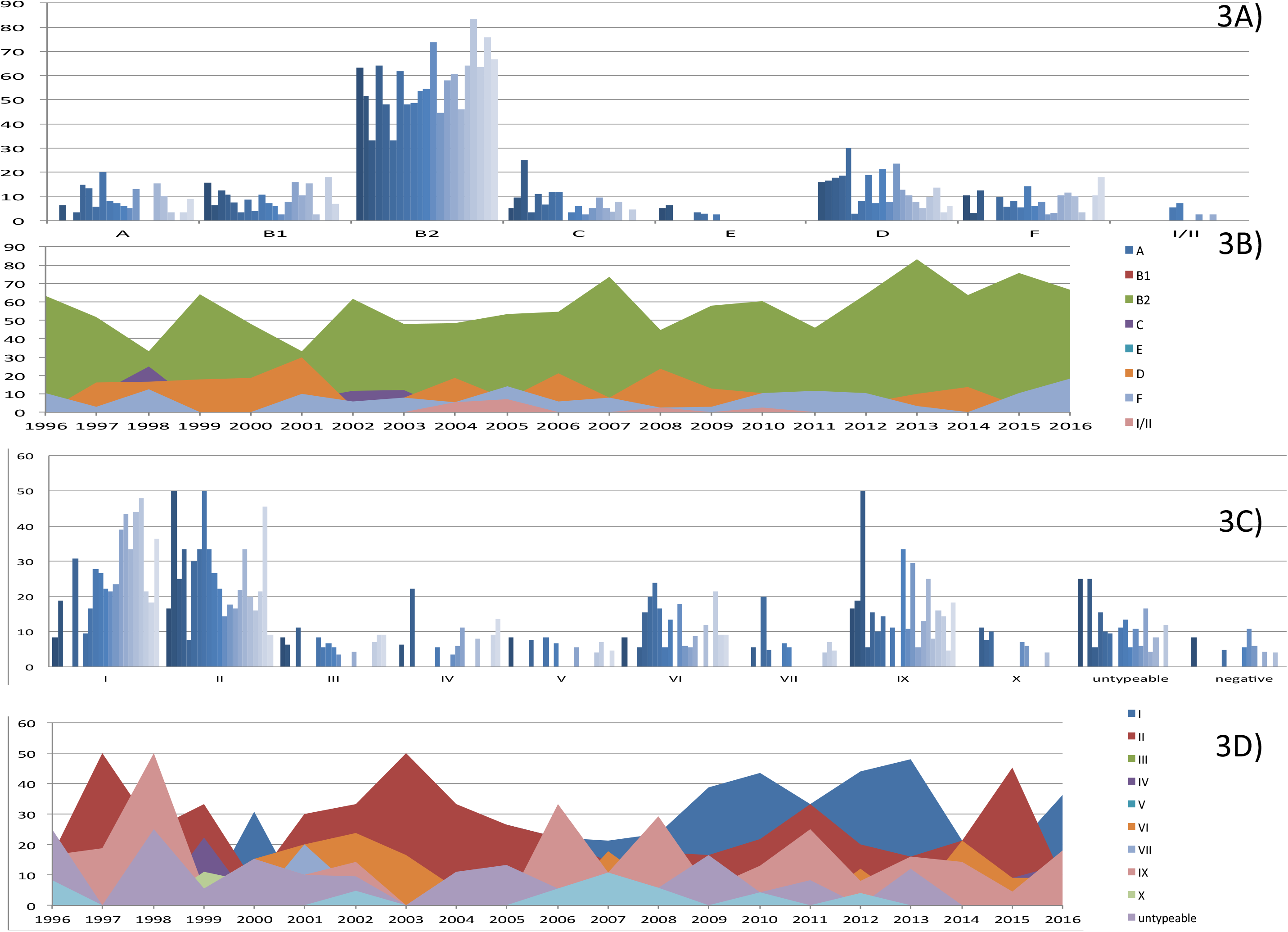
Temporal Distribution of *E. coli* populations at Hospital Universitario Ramón y Cajal, HURyC (1996-2016). Panels A and B) Temporal distribution of *E. coli* phylogroups; panels C and D) Temporal distribution of *E. coli* B2 subgroups.

Almost half of patients with STc131 (46%) and all predominant B2-II (STc73), B2-III (STc127), B2-VI (STc12), B2-IX (STc95) persistent clones were admitted to the medical emergency ward with an established BSI, suggesting the frequent presence of these clones in the community. We also detected the same PFGE patterns in isolates from different patients for *E. coli* of B2-I-ST131 (clades H22, C1, C2), B2-II (ST73), B2-III (ST127), B2-Iv, B2-VI (ST12), B2-IX (ST95) (Table S1).

#### Temporal variation

Except for phylogroups B2-I (ST131) and phylogroup D, the trends of the phylogroups did not significantly change during the period analyzed (Figure 3). However, the occurrence of major subgroups (B2-II, B2-IX, B2-VI, B-IV, B2-VII) greatly varied through the years suggesting episodes of transmission with transient amplifications.

The STc131 *E. coli* was the only group increasingly recovered coinciding with its global amplification (24). The ST131 clade B was initially detected in 1996, and remained steadily identified since then. In the current study, the ST131 of clade A was first detected in 2004, but we had identified STc131 clade A in clinical isolates of TEM-4 and TEM-24 producers from 1991 and 2000, respectively, in other studies of the group (25, 26) For the predominant ST131 clade C, the subclade C0 (H30, FQ^S^) was detected in 2000, followed by C1 (H30-R, FQ^R^) in 2004, C2 (H30-Rx, FQ^R^ *bla*_CTX-M-15_) in 2006, and C1-M27 (H30-Rx, FQ^R^ *bla*_CTX-M-27_) in 2016 (Figure 4). Further analysis of PFGE and WGS revealed clonal amplifications (see next section).

**Figure 4.**
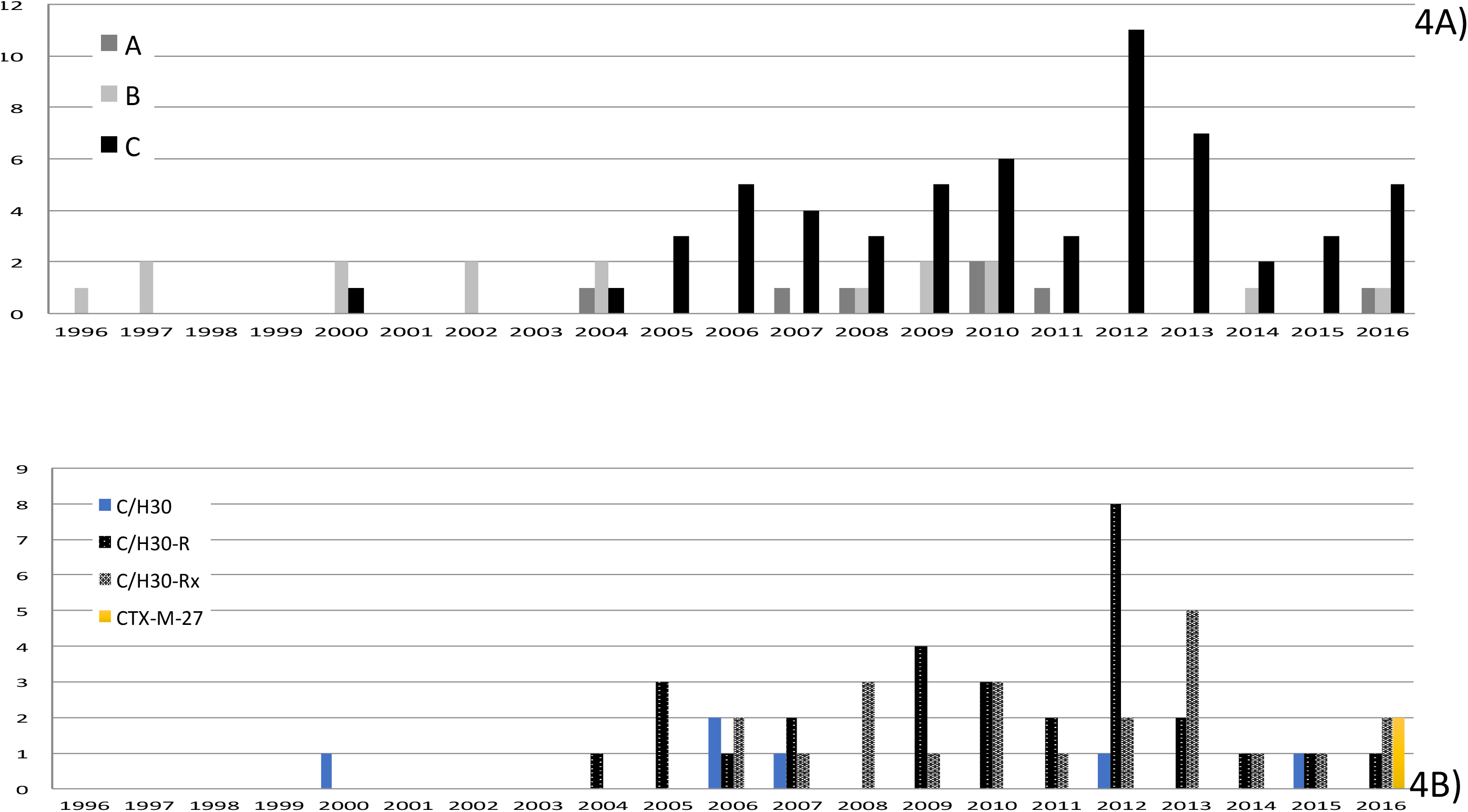
Temporal Distribution of *E. coli* populations at HRyC (1996-2016). (A) Distribution of STc131 clades; (B) Timeline of the STc131 subclades C0, C1.

#### Distribution by Age

All phylogenetic groups were present in patients of all age groups, although differences were observed for subgroups B2-I (STc131), B2-II, B2-VI, and B2-IX. This distribution implies the clear predominance of B2-I in the elderly (>80 years) and B2-II in individuals younger than 45 years. Others showed a bimodal distribution as B2-IX and B2-VI (Figure S2).

### Antimicrobial susceptibility

Figure S3 shows the antimicrobial susceptibility patterns in the sample of 649 *E. coli* isolates, revealing that ciprofloxacin (Cip) resistance (Cip^R^) appears in between 35% and 50% of the isolates of each phylogroup. Although the lowest Cip^R^ rate corresponded to B2 phylogroup (17%), it is of note that most Cip^R^ B2 isolates were STc131 (70% vs. 3.5% in non-ST131 B2, with major B2 subgroups II, III, IV, VI and IX being susceptible). Only 6.1% of the total number of B2 isolates showed a 3GC^R^ phenotype compatible with the production of an extended-spectrum beta-lactamase (ESBL), further identified as CTX-M-15, CTX-M-14, and CTX-M-27, and was only detected among STc131 isolates. Resistance to ampicillin (70.4%), streptomycin (39.6%), nalidixic acid (32.8%), and cotrimoxazole (45%–50%) was frequent among the isolates of the various phylogroups. Remarkable differences were observed for amoxicillin-clavulanic acid, kanamycin, gentamicin, tetracycline, and chloramphenicol, mostly due to the phylogroup C isolates, all clonally unrelated (data not shown).. Susceptibility to the 13 antimicrobials tested was observed in 17% of the total number of isolates tested, with the frequency of susceptible isolates higher among those of phylogroups A, B1, B2, and F (15% each) than those of phylogroups D (10%), E (1%), and C (0%). Within B2, a pan-susceptibility pattern was more frequent for non-STc131 than for STc131 isolates (26.8% vs. 3.7%). The B2-STc131 strains showed a multiresistant profile (resistance against ≥ antimicrobial agents of different families) more frequently than other B2 members (56.6% vs. 11.6%; p <.001).

### Virulence-associated gene profiling in the B2 phylogroup

According to the content of the VAG genes, the B2 *E. coli* strains were clustered into 2 large groups and 10 subgroups, each comprising numerous gene combinations and showing variable redundancies (Figure 5). While more than half (56%) of the B2-I (STc131) strains clustered in VAG 1 group, with predominance of the subcluster VAG 1.5, *E. coli* isolates of B2-II, B2-III, B2-V, B2-VII, B2-IX, and B2-X (54-75%) clustered in the VAG 2 group and more specifically, in the VAG subgroups 2.6, 2.2, 2.9, 2.6, 2.7, and 2.4, respectively (ordered by frequency). Despite some VAG variability within each B2 phylogenetic subgroups, the results suggest a conserved genetic structure related to virulence and colonization in the B2 strains analyzed in this study, in part due to the presence of some persistent (“epidemic”) strains identified through the years. A detailed analysis of virulence content is provided in Supplementary Text and Figures S4, S5, and S6.

**Figure 5.**
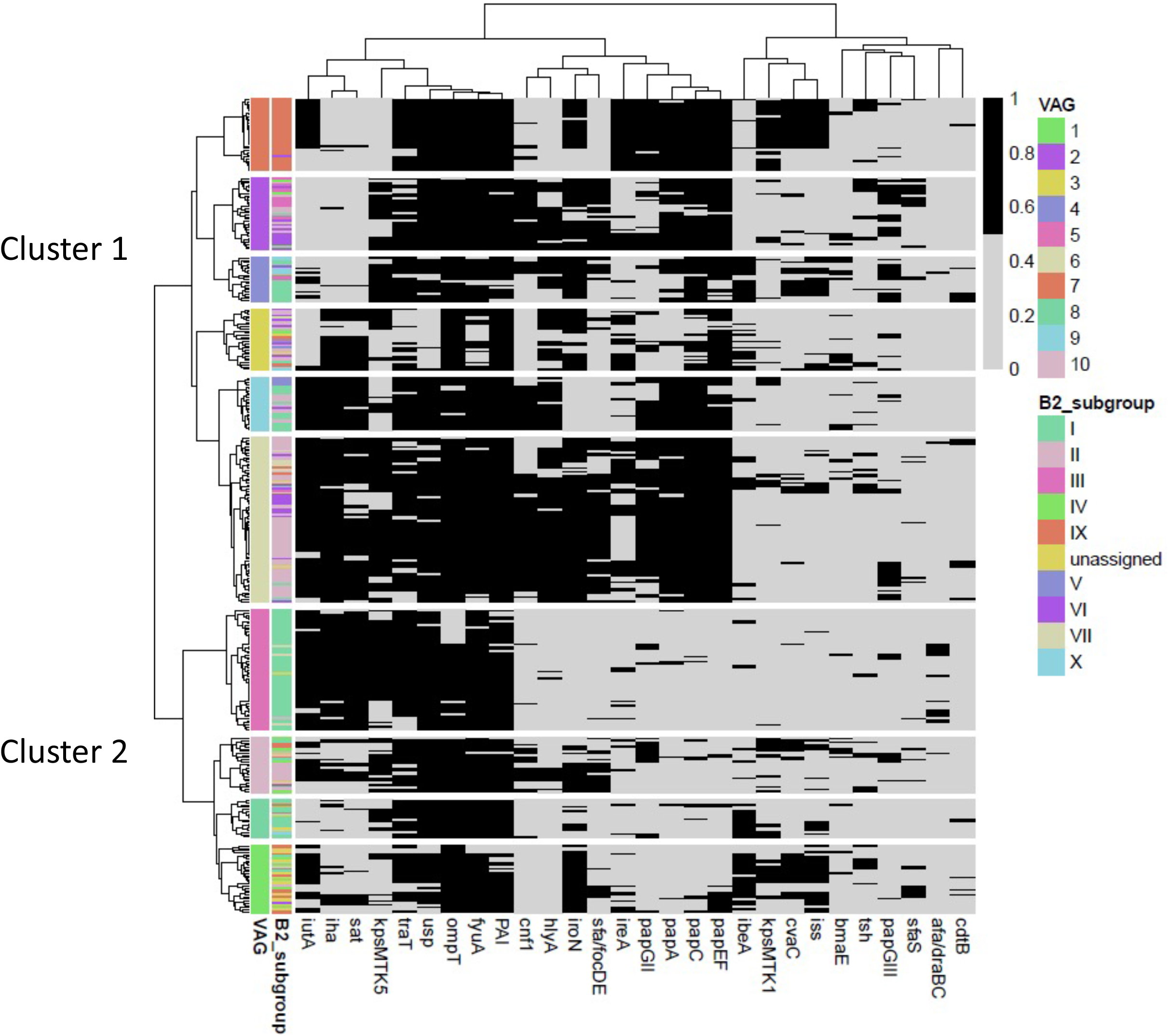
Heatmap of 29 virulence factors (presence/absence) for the 348 B2 phylogenetic group strains causing BSI at Ramón y Cajal University Hospital (1996–2016). VAGs: Virulence Associated groups. VAGs were determined by hierarchical clustering using the Ward method and Jaccard similarity distance. Abbreviations of the VAGs tested. *Toxins*: *hlyA* (α-Hemolysin), *sat* (secreted autotransporter toxin, a serine protease), *cnf1* (cytotoxic necrotizing factor 1), *cdtB* (cytolethal distending toxin), *tsh* (temperature-sensitive hemagglutinin); *siderophores*: *iroN* (salmochelin receptor)*, iutA* (aerobactin synthesis, receptor)*, ireA* (iron-regulated element, catecholate siderophore)*, fyuA* (yersiniabactin receptor); *adhesins*: fimbriae type P (*pap*GI, *pap*GII, *pap*GIII, *pap*A, *pap*C, *pap*EF), *sfa/focDE* (type S fimbriae, sfa/foc S and F1C fimbriae); *afa/draBC*, Adhesinsd afa/dra Dr antigen-binding adhesins (AFA I-III, Dr, F1845; *bmaE* (blood group M-specific adhesin*; iha* (iron-regulated-gene-homologue adhesin); *protectins*: kpsMT II (group II capsule synthesis, e.g., K1, K5, K12). kpsMT III Group III capsule synthesis (e.g., K3, K10, K54), *traT* (surface exclusion; serum resistance-associated), *invasins*: ibeA-C (invasion of brain endothelium IbeA); *miscellanea*: *cvaC* (microcin/colicin V; on plasmids with *traT, iss, iuc/iut*), *ompT* (outer membrane protein T), *usp* (uropathogenic-specific protein, bacteriocin), PAI (malX, a PAI CFT073 marker), *iss* (increased serum survival, outer membrane protein).

#### Clonal relationships

PFGE typing of isolates of B2 subgroups II-X revealed highly/identical patterns of XbaI-digested genomic DNA for isolates within B2 subgroups II (STc73), IX (STc95), VI (ST12), and III (STc127) collected during variable long periods of time (2-17 years) (Table S1). Analysis of pairwise SNP distances between isolates and the tree topology, suggest the presence of different lineages of different origin for each STs, which showed identical or highly similar Xba-I digested genomic DNA patterns. It is of note that such lineages/PFGE types differ in a variable number of SNPs (18-94SNPs, some individual isolates of each cluster differing in close 100SNPs) (Figures 6A and 6B). Although genetic distances estimated as SNP constitutes the initial basis to determine the similarity between isolates, available thresholds for establishing such similarity are mostly based on outbreak investigations and conventional mutation rates, the interpretation of SNP/ allelic values to conclude on the similarity of isolates remaining challenging. Nonetheless, the similarity of the PFGE profiles and the time lapse between isolates suggests the successful and long-lasting transmission of certain lineages in the community.

**Figure 6.**
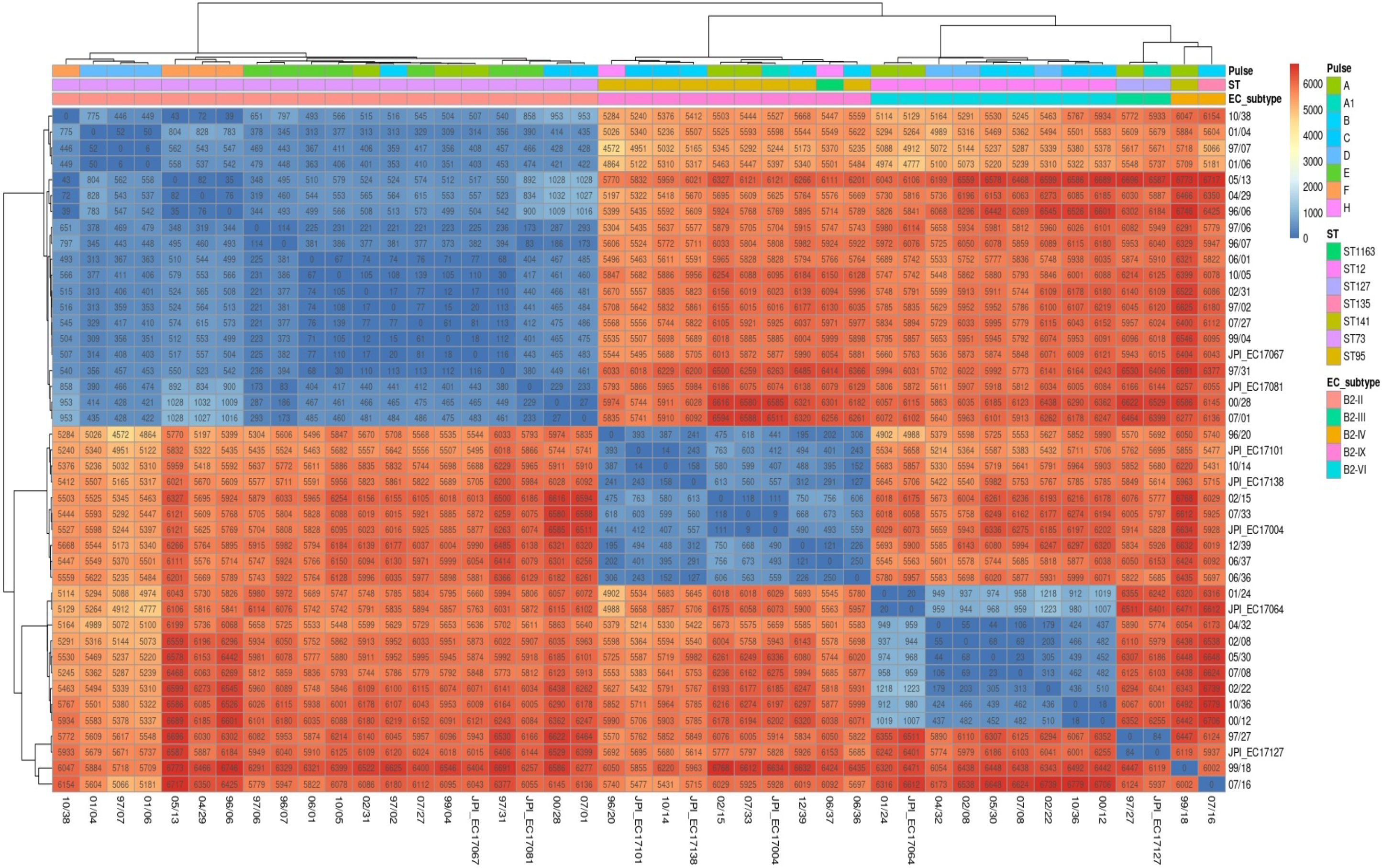
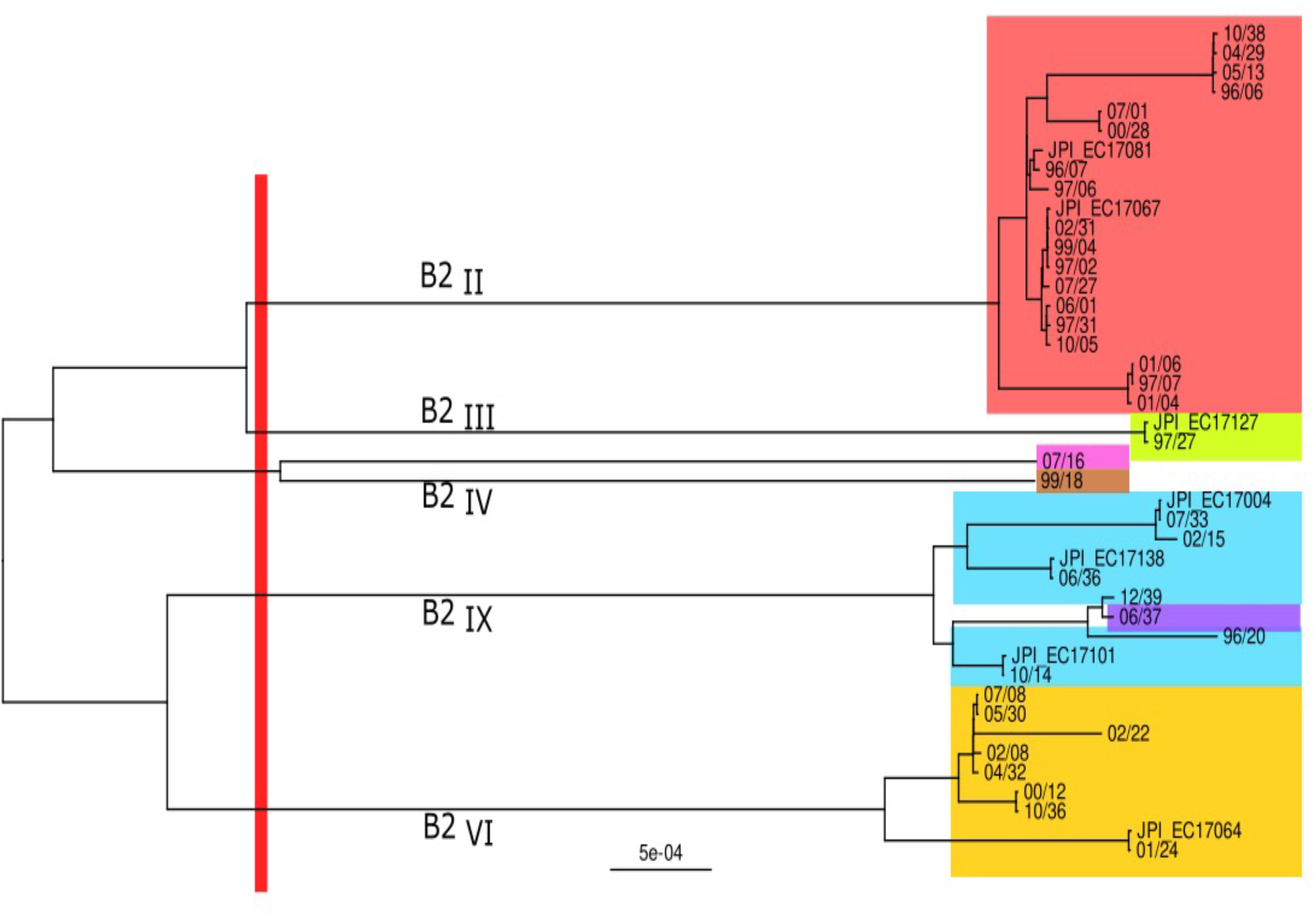
Characterization of B2 E. coli strains fully sequenced. Panel 6A). Heatmap of pairwise SNP distances based on the absolute differences of SNPs. The colors of the boxes identify the different SNP clades (B2_II_-ST73, B2_IX_-STc95, B2_IV_-ST12, B2_III_-ST127). Panel B) Maximum-likelihood phylogenetic tree of the whole genome performed using FastTree (60). Clades are named and colored according to the ST subgroup:: B2_II_-ST73 (red), B2_IX_-ST95 (blue) and ST1163 (violet), B2_III_-ST127 (pink), B2_VI_-ST12 (yellow). SNP numbers represent the range of individual comparisons.

#### Gene Accession numbers

The sequences of the genomes have been registered in the BioProject database with the reference PRJNA775650. The references for the BioSample accessions are SAMN22610064-SAMN22610106.

## DISCUSSION

This work, one of the very few long-term longitudinal studies -and the first in Spain-to analyse the dynamics of *E. coli* phylogroups involved in BSIs for a long period of time at a single centre, reveals intertwined waves of CA+HCA-BSI and HA-BSI episodes caused by strains of both B2 and non-B2 phylogroups for decades. This would have occurred until BSI episodes caused by B2-populations of *E. coli* overtook those of non-B2 and the B2-I-STc131 became transiently predominant and predate new waves of other major B2 subgroups. Such dynamics is similar to that observed in the UK (17) but our study further suggests that the burden of CA-BSI and HA-BSI infections are closely related, genomic and epidemiological data suggesting differences between the evolutionary pathways of different *E. coli* phylogroups and B2 subgroups.

B2-I (STc131) was the only lineage that significantly increased during the period analyzed, and this transient increase was due to the expansion of both C1 and C2 subclades (mostly ESBL negative), and probably, to CTX-M-27 in the last years, according to the emergence of this subclade in the community after the end of this study (23). Although the epidemiology of the STc131 has extensively been documented (11), this work and that by Kallonen (17) are the only ones that reveals the coexistence of all STc131 subclades which, after a transient amplification, are incorporated into the *E. coli* population structure. The other predominant B2 subgroups varied greatly through the period studied, with some peaks of B2-II (STc73), B2-IX (STc95), B2-VI (STc12) clones and also others (e.g. B2-VII, B2-X, non-I/II) which are poorly documented in the literature. The identification of isolates of each predominant B2 phylogroups/STs, mostly collected from patients at the hospital emergency wards through a wide range of years (2-17 years), only differing in tens of SNPs (33-94 SNPs) and showing identical or highly similar PFGE types, is of utmost relevance and suggests a reservoir of highly stable “epidemic” strains in the community that would have evolved within or through different individuals or groups of related individuals (28). Note that the interpretation of SNP/ allelic values to conclude on the similarity of isolates is still challenging because the thresholds currently used are based on outbreak investigations and reported mutation rates under laboratory conditions, and thus, only extrapolated to isolates collected in short period of times.

Analysis of the population structure of commensal *E. coli* by Massot et al demonstrated a shift In Western countries from phylogroups A and D in 1970s to B2 and F in 2000s which could be influenced by the increasing use of antibiotics (9). However, the steady increase of *E. coli* resistant to Cip and 3GCs observed since the early 1990s occurs in non-B2 groups and B2-ST131 subclade C and contrasts with the antibiotic susceptibility to Cip, 3GC and other drugs of most predominant STs of B2-nonST131 isolates and ST131 subclades A and C0 (this study,(29). Besides antibiotic pressure, significant changes in healthcare delivery and in the human population structure determinants (age, interconnectedness, diet) have recently been suggested to explain the amplification of particular strains (36–38) but any of these factors do not fully justify the results. Cip^R^ has recently been considered an advantage for the fitness of high-risk clones of *E. coli* as ST131 subclade C and also of other bacterial species (39, 40). Some Cip ^R^ clones are predominant in the elderly, with a sustained pattern in hospital (or day-care-center) admissions and personal clinical history that facilitates the acquisition of other antibiotic resistances (41). An increasing cumulative “elderly microbiota reservoir” of antimicrobial-resistant subpopulations of various bacterial species would be congruent with a high occurrence of CA+HCA BSI episodes at admission, with the age of the patients in our sample, and with the identification of identical PFGE types in groups of unrelated patients. However, the detection of B2 isolates of other non-ST131 clade C groups in people of all ages but predominantly in the range of 15-45y and also in other non-human hosts (29, 36–39), eventually being able to circulate between them (28, 36), would reinforce that they are part of the normal microbiota. Note than the notion of “pandemic” implies the extended spread of the clones in the community, and the increase in number and exposure to different hosts and environments necessarily tends to an increase in genetic diversity (11). In fact, the notion that the “high-risk clones” emerge from the epidemicity of commensals which precedes the spread of multi-resistant bacterial clones (36, 40).

The correlation between VAG profiles and B2 subgroups reflects an apparent structure of B2 populations in agreement with the ecological niche specialization theory, considering that the niche of a population is the result of the niches occupied by all its individuals (7, 41–45), so that the niche is evolving itself (46, 47). Although the concordance of specific VAGs with isolates of B2-II and B2-IX subgroups is congruent with their classical categorization as “uropathogenic” *E. coli*, such a way explaining their bimodal distribution in different age groups, most B2-I strains (STc131 clade C) exhibit a low number of classical EC-VAGs (48, 49). To explain the expansion of the STc131 and other emerging lineages, such as STc648 (phylogroup F), studies that have applied mathematical models to a high number of genomes suggest that negative frequency-dependent selection of previously rare populations might have favored their increase (14, 49), and will foster the expansion of some others in the future. However, these studies are fully focused on the microorganisms and do not allow us to associate the observed changes with the type of hosts located either in hospital or community compartments.

We are aware that the size of the sample analyzed, despite of the high number of carefully randomized isolates analyzed, only represents the 10% of the total number of BSI in our institution during the studied period. However, the long period analyzed at a single institution helps identifying relevant epidemiological conditions that illustrates the expansion or particular *E. coli* populations observed in our and other areas during recent years. Currently, the B2 *E. coli* strains are relevant units of BSI pathogenicity, which should correlate with their success in particular microecological landscapes, in part determined by recent interventions exerted on particularly fragile human populations (mostly elderly patients), and also because the cumulative effect of interventions during lifetime. This view is in agreement with the concept of incorporating ecological features in the identification of the fundamental units of bacterial diversity (38) and pathogenicity. In fact, particular clones and lineages are differentially represented in a “human microbiota reservoir” flowing from the community to the hospital and vice versa, where they can either be selected or coexist as predicted by an evolutionarily stable strategy (38). The evolutionary trajectories of recently amplified major lineages indicate the relevance of undetected selective events, which could be further amplified by the acquisition of antimicrobial resistance in settings under (or not) antibiotic pressure (37, 38, 50). The early detection of the abundance and diversity of B2 subgroups of clinical significance for BSI in metagenomic samples of community-based individuals and the analysis of common causes that enhance their selection, either in the hospital, nursing homes or the community, are priority research challenges that warrant attention in an era that favor the pandemics of microorganisms and antimicrobial resistance (51).

## METHODS

### Study design

Ramón y Cajal University Hospital is a tertiary-level public health center with 1155 beds that provides attention to 600,000 habitants in the Northern area of Madrid (Spain), which reflects a pyramid-age of “declining type”, has full accessibility to primary attention care and have a predominant medium-high socioeconomic level. Of the total 21,695 positive blood cultures detected between January 1996 and December 2016, we identified 7165 *E. coli* that represented 1 isolate per patient and per BSI episode. A sample of nearly 10% of this *E. coli* collection, stratified by sex, age, and antimicrobial resistance pattern was sorted by statistical randomization (Stata Statistical Software: Release 17. College Station, TX: StataCorp LLC) and was further analyzed (649 *E. coli* isolates from 339 females and 310 males; < 1-98 years of age). The study was approved by the Ethics Committee of our hospital.

BSIs are classified as hospital acquired (HA), community acquired (CA), and community-onset healthcare-associated (HCA), according to the date of the sample collection after patient admission and the patient exposure to hospitals before the BSI episode (19, 50). Due to the inaccessibility to all the medical records of patients enrolled in this study, and their advanced age, we classified the episodes into HA (if the blood culture was obtained at the ICU, surgical, or medical areas after 48 h of admission) and CA+HCA (if the blood culture was obtained at the hospital emergency wards or at the day care centers) categories. UTIs were considered as the origin of BSI if *E. coli* was recovered from both the urine and blood samples, with a difference of ±24 h.

### Characterization of the bacterial isolates

Blood culture isolates of *E. coli* are routinely frozen and stocked in skimmed milk at −70°C, and were subcultured onto brain-heart infusion agar prior to analysis. Bacterial susceptibility against 13 antibiotics (ampicillin, amoxicillin-clavulanic acid, cefotaxime, ceftazidime, meropenem, nalidixic acid, ciprofloxacin, streptomycin, kanamycin, gentamicin, tetracycline, chloramphenicol, and trimethoprim/sulfamethoxazole) was performed by the disk-diffusion method (52).

Multiplex PCR assays allowed classifying the *E. coli* isolates into major phylogenetic groups A, B1, B2, C, D, E, and F (53); B2 subgroups (B2 I-X) (54); and B2-I-STc131 *E. coli* serotypes (O16/O25b), clades (H41, H22, H30), and the H30 subclades C0 [H30, fluoroquinolone susceptible (FQ^S^)], C1 [H30-R, fluoroquinolone resistant (FQ^R^)], C2 (H30Rx, FQ^R^+*bla*_CTX-M-15_), and C1-M27 (H30-Rx, FQ^R^+ *bla*_CTX-M-27_) (54, 55). STc131 isolates were further analyzed by pulsed field gel electrophoresis (PFGE). The presence of 29 virulence-associated genes (VAGs) was determined for all B2 isolates by PCR (6, 56).

Clonal relationship between B2 isolates was preliminary established by Pulsed field gel electrophoresis (PFGE) according the PulseNet website (https://pulsenetinternational.org/protocols/pfge/). Isolates showing the same or highly related PFGE profiles (1-3 bands) were selected for WGS (Table 1).

### Whole genome sequencing

Forty-five solates corresponding to isolates showing similar or highly related PFGE types were selected for WGS. DNA was extracted from 5 mL of overnight cultures using the Wizard Genomic DNA Purification Kit (Promega Corp., Madison, WI, USA) and DNA concentration was measured using a Qubit™ Fluorometer and Nanodrop 2000 (Thermo Scientific, Waltham, MA, USA). WGS was performed using the Illumina-NovaSeq 6000 platform (Oxford Genome Center, Wellcome Centre for Human Genetics Oxford, UK) with 2% 150 bp paired-end reads. Quality control and sequence filtering was done using the FastQC v.0.11.8 (https://www.bioinformatics.babraham.ac.uk/projects/fastqc/) and Prinseq-lite-0.20.3 (http://prinseq.sourceforge.net/) tools, respectively.

The paired-end reads were *de novo* assembled using SPAdes v.3.14.1 (56) and then, they were annotated with Prokka (57) The phylogenetic analysis was obtained using PATO (58) and fasttree2.1 (59). The heatmap resulted from a whole genome snp pairwise comparison (PATO) and the tree was made with ggtree (60). *In silico* MLST assignment was performed using MLST v2.16.1 (https://github.com/tseemann/mlst).

### Statistical analysis

To calculate statistical significance, the chi-squared test, a 2-sample t-test for normally distributed variables, and Kendall’s correlation were used, considering p-values <0.05 to be statistically significant

## AUTHORS’ CONTRIBUTIONS

I.R. performed the wetlab work of the isolates collected from 1996–2012, assisted by C.R., and participated in the early study design, the data analysis, and the very initial draft of the manuscript. A.S.F and M.S, performed the wet lab work of the isolates collected from 2012– 2016, the categorization of all ST131s included in the study, the data analysis, and the revision of the manuscript. SAG performed the PFGE analysis and its comparative analysis with WGS data of the isolates. MDFdB, CB, and V.F.L performed all the bioinformatic and statistical analysis of the results and the revision of the manuscript. J.Z participate in the sample design and statistical analysis. E.L and RC provided valuable information from the clinical microbiology lab, patient records, and revised the manuscript. PM contribute in the antibiotic susceptibility analysis. FB joined the study design, revise the different versions of the paper, and made relevant contributions to the discussion. T.M.C. participated in the study design, the data analysis, and wrote the manuscript. All the authors have read and approved the final version of this document.

## SUPPLEMENTARY FIGURE CAPTIONS

**Supplementary Text.** Virulence-associated gene profiling in the B2 phylogroup of *Escherichia coli*

**Table S1. Relationship between predominant clonal lineages isolated in HURyC (1996-2016)**. The table reflects the Sequence types (STs), PFGE profiles, date of isolation and hospital ward where the blood culture was taken.

**Supplementary Figure S1. Distribution of *E. coli* phylogroups and B2 phylogenetic subgroups.** (A) *E. coli* phylogroups; (B) *E. coli* B2 phylogenetic subgroups; (C) *E. coli* B2_I_ (STc131) clades A, B, C, and subclades C0/H30 (FQ^S^), C1/H30-R (FQ^R^), C2/H30Rx (FQ^R^+*bla*_CTX-M-15_), and C1-M27/H30-R (FQ^R^+ *bla*_CTX-M-27_).

**Supplementary Figure S2. Diversity of *E. coli* per age group.** (A) Per phylogroup; (B) per B2 subgroup. Half of the B2 isolates corresponded to B2_II_ and B2_IX_ (33% and 21%) for patients between 15 and 45 years, B2-I and B2-II (23% and 25%–28%) for patients between 45 and 80 years, and B2_I_ and B2_IX_ (36% and 20%) for patients >80 years.

**Supplementary Figure S3. Antibiotic susceptibility of B2-*Escherichia coli* causing BSI.** Abbreviations: AMC, amoxicillin-clavulanic acid; AMP, ampicillin; CAZ, ceftazidime; CHL, chloramphenicol; CIP, ciprofloxacin; CTX, cefotaxime; GEN, gentamicin; KAN, kanamycin; MEM, meropenem; NAL, nalidixic acid; STR, streptomycin; TET, tetracycline; TMP/SXT, trimethoprim-sulfamethoxazole; Susceptible, susceptible to all 13 antibiotics analyzed.

**Supplementary Figure S4. Heatmap displaying the relative abundance of B2 phylogenetic groups and VAG profiles**. Variables showing goodness-of-fit are shown in red, whereas dark blue regions indicate poor frequencies. Clustering of the B2 subgroups and VAGs and the assembly statistics were performed using a hierarchical clustering method in R (hclust). Abbreviations of the VAGs tested. *Toxins*: *hlyA* (α-Hemolysin), *sat* (secreted autotransporter toxin, a serine protease), *cnf1* (cytotoxic necrotizing factor 1), *cdtB* (cytolethal distending toxin), *tsh* (temperature-sensitive hemagglutinin); *siderophores*: *iroN* (salmochelin receptor)*, iutA* (aerobactin synthesis, receptor)*, ireA* (iron-regulated element, catecholate siderophore)*, fyuA* (yersiniabactin receptor); *adhesins*: fimbriae type P (*pap*GI, *pap*GII, *pap*GIII, *pap*A, *pap*C, *pap*EF), *sfa/focDE* (type S fimbriae, sfa/foc S and F1C fimbriae); *afa/draBC*, Adhesinsd afa/dra Dr antigen-binding adhesins (AFA I-III, Dr, F1845; *bmaE* (blood group M-specific adhesin*; iha* (iron-regulated-gene-homologue adhesin); *protectins*: kpsMT II (group II capsule synthesis, e.g., K1, K5, K12). kpsMT III Group III capsule synthesis (e.g., K3, K10, K54), *traT* (surface exclusion; serum resistance-associated), *invasins*: ibeA-C (invasion of brain endothelium IbeA); *miscellanea*: *cvaC* (microcin/colicin V; on plasmids with *traT, iss, iuc/iut*), *ompT* (outer membrane protein T), *usp* (uropathogenic-specific protein, bacteriocin), PAI (malX, a PAI CFT073 marker), *iss* (increased serum survival, outer membrane protein).

**Supplementary Figure S5. Heatmap displaying the relative abundance of B2 phylogenetic groups and VAGs**. Variables showing goodness-of-fit are shown in red, whereas dark blue regions indicate poor performance. Clustering of the B2 subgroups and VAGs and the assembly statistics were performed using a hierarchical clustering method in R (hclust).

Abbreviations of the VAGs tested. Genes coding for toxins: *hlyA* (α-Hemolysin), *sat* (secreted autotransporter toxin, a serine protease), *cnf1* (cytotoxic necrotizing factor 1), *cdtB* (cytolethal distending toxin), *tsh* (temperature-sensitive hemagglutinin); *siderophores*: *iroN* (salmochelin receptor)*, iutA* (aerobactin synthesis, receptor)*, ireA* (iron-regulated element, catecholate siderophore)*, fyuA* (yersiniabactin receptor). Genes coding for *adhesins*: fimbriae type P (*pap*GI, *pap*GII, *pap*GIII, *pap*A, *pap*C, *pap*EF), *sfa/focDE* (type S fimbriae, sfa/foc S and F1C fimbriae); *afa/draBC*, Adhesinsd afa/dra Dr antigen-binding adhesins (AFA I-III, Dr, F1845; *bmaE* (blood group M-specific adhesin*; iha* (iron-regulated-gene-homologue adhesin); *protectins*: kpsMT II (group II capsule synthesis, e.g., K1, K5, K12). *kpsMT* III Group III capsule synthesis (e.g., K3, K10, K54), *traT* (surface exclusion; serum resistance-associated). Genes coding for *invasins*: *ibe*A-C (invasion of brain endothelium IbeA). Genes coding for other genes: *cvaC* (microcin/colicin V; on plasmids with *traT, iss, iuc/iut*), *ompT* (outer membrane protein T), *usp* (uropathogenic-specific protein, bacteriocin), PAI (*malX*, a PAI CFT073 marker), *iss* (increased serum survival, outer membrane protein).

**Supplementary Figure S6. Heatmap the relative abundance of each VAG in each VAG group.** Variables showing goodness-of-fit.

Abbreviations of the VAGs tested. Genes coding for toxins: *hlyA* (α-Hemolysin), *sat* (secreted autotransporter toxin, a serine protease), *cnf1* (cytotoxic necrotizing factor 1), *cdtB* (cytolethal distending toxin), *tsh* (temperature-sensitive hemagglutinin); *siderophores*: *iroN* (salmochelin receptor)*, iutA* (aerobactin synthesis, receptor)*, ireA* (iron-regulated element, catecholate siderophore)*, fyuA* (yersiniabactin receptor). Genes coding for *adhesins*: fimbriae type P (*pap*GI, *pap*GII, *pap*GIII, *pap*A, *pap*C, *pap*EF), *sfa/focDE* (type S fimbriae, sfa/foc S and F1C fimbriae); *afa/draBC*, Adhesinsd afa/dra Dr antigen-binding adhesins (AFA I-III, Dr, F1845; *bmaE* (blood group M-specific adhesin*; iha* (iron-regulated-gene-homologue adhesin); *protectins*: kpsMT II (group II capsule synthesis, e.g., K1, K5, K12). *kpsMT* III Group III capsule synthesis (e.g., K3, K10, K54), *traT* (surface exclusion; serum resistance-associated). Genes coding for *invasins*: *ibe*A-C (invasion of brain endothelium IbeA). Genes coding for other genes: *cvaC* (microcin/colicin V; on plasmids with *traT, iss, iuc/iut*), *ompT* (outer membrane protein T), *usp* (uropathogenic-specific protein, bacteriocin), PAI (*malX*, a PAI CFT073 marker), *iss* (increased serum survival, outer membrane protein).

**Supplementary Figure S7. Heatmap of 29 VAGs (presence/absence)**. The heatmap was annotated with the FimH allele, ST131 serotype, patient age, and sex.

Abbreviations of the VAGs tested. Genes coding for toxins: *hlyA* (α-Hemolysin), *sat* (secreted autotransporter toxin, a serine protease), *cnf1* (cytotoxic necrotizing factor 1), *cdtB* (cytolethal distending toxin), *tsh* (temperature-sensitive hemagglutinin); *siderophores*: *iroN* (salmochelin receptor)*, iutA* (aerobactin synthesis, receptor)*, ireA* (iron-regulated element, catecholate siderophore)*, fyuA* (yersiniabactin receptor). Genes coding for *adhesins*: fimbriae type P (*pap*GI, *pap*GII, *pap*GIII, *pap*A, *pap*C, *pap*EF), *sfa/focDE* (type S fimbriae, sfa/foc S and F1C fimbriae); *afa/draBC*, Adhesinsd afa/dra Dr antigen-binding adhesins (AFA I-III, Dr, F1845; *bmaE* (blood group M-specific adhesin*; iha* (iron-regulated-gene-homologue adhesin); *protectins*: kpsMT II (group II capsule synthesis, e.g., K1, K5, K12). *kpsMT* III Group III capsule synthesis (e.g., K3, K10, K54), *traT* (surface exclusion; serum resistance-associated). Genes coding for *invasins*: *ibe*A-C (invasion of brain endothelium IbeA). Genes coding for other genes: *cvaC* (microcin/colicin V; on plasmids with *traT, iss, iuc/iut*), *ompT* (outer membrane protein T), *usp* (uropathogenic-specific protein, bacteriocin), PAI (*malX*, a PAI CFT073 marker), *iss* (increased serum survival, outer membrane protein).

## References

1. de Kraker MEA, Jarlier V, Monen JCM, Heuer OE, van de Sande N, Grundmann H. 2013. The changing epidemiology of bacteraemias in Europe: trends from the European Antimicrobial Resistance Surveillance System. Clin Microb Infect 19:860–8.

2. Jarlier V, Diaz Högberg L, Heuer OE, Campos J, Eckmanns T, Giske CG, Grundmann H, Johnson AP, Kahlmeter G, Monen J, Pantosti A, Rossolini GM, van de Sande-Bruinsma N, Vatopoulos A, Żicka D, Žemličková H, Monnet DL, Simonsen GS, Ears-Net Participants. 2019. Strong correlation between the rates of intrinsically antibiotic-resistant species and the rates of acquired resistance in Gram-negative species causing bacteraemia, EU/EEA, 2016. Euro Surveill 24(33):1800538.

3. Laupland KB, Gregson DB, Church DL, Ross T, Pitout JDD. 2008. Incidence, risk factors and outcomes of *Escherichia coli* bloodstream infections in a large Canadian region. Clin Microb Infect 14:1041–1047.

4. Bou-Antoun S, Davies J, Guy R, Johnson AP, Sheridan EA, Hope RJ. 2016. Descriptive epidemiology of *Escherichia coli* bacteraemia in England, April 2012 to March 2014. Euro Surveill 21(35):30329.

5. Tenaillon O, Skurnik D, Picard B, Denamur E. 2010. The population genetics of commensal *Escherichia coli*. Nat Rev Microbiol 8:207–17.

6. Johnson JR, Russo TA. 2018. Molecular Epidemiology of Extraintestinal Pathogenic *Escherichia coli*. EcoSal Plus 1(1).

7. Le Gall T, Clermont O, Gouriou S, Picard B, Nassif X, Denamur E, Tenaillon O. 2007. Extraintestinal virulence is a coincidental by-product of commensalism in B2 phylogenetic group *Escherichia coli* strains. Mol Biol Evol 24:2373–84.

8. Nowrouzian FL, Clermont O, Edin M, Östblom A, Denamur E, Wold AE, Adlerberth I. 2019. *Escherichia coli* phylogenetic B2 subgroups in the infant gut microbiota: predominance of uropathogenic lineages in Swedish infants and enteropathogenic lineages in Pakistani infants. Appl Environ Microbiol 85:e01681–19.

9. Massot M, Daubié A-S, Clermont O, Jauréguy F, Couffignal C, Dahbi G, Mora A, Blanco J, Branger C, Mentré F, Eddi A, Picard B, Denamur E, The Coliville Group. 2016. Phylogenetic, virulence and antibiotic resistance characteristics of commensal strain populations of *Escherichia coli* from community subjects in the Paris area in 2010 and evolution over 30 years. Microbiology 162:642–650.

10. Olesen B, Scheutz F, Menard M, Skov MN, Kolmos HJ, Kuskowski MA, Johnson JR. 2009. Three-decade epidemiological analysis of *Escherichia coli* O15:K52:H1. J Clin Microbiol 47:1857–1862.

11. Baquero F, Martínez JL, F. Lanza V, Rodríguez-Beltrán J, Galán JC, San Millán A, Cantón R, Coque TM. 2021. Evolutionary Pathways and Trajectories in Antibiotic Resistance. Clin Microbiol Rev 34:e0005019.

12. Manges AR, Geum HM, Guo A, Edens TJ, Fibke CD, Pitout JDD. 2019. Global Extraintestinal Pathogenic *Escherichia coli* (ExPEC) Lineages. Clin Microbiol Rev 32:e00135–18.

13. Alhashash F, Weston V, Diggle M, McNally A. 2013. Multidrug-resistant *Escherichia coli* bacteremia. Emerg Infect Dis 19:1699–1701.

14. Schaufler K, Semmler T, Wieler LH, Trott DJ, Pitout J, Peirano G, Bonnedahl J, Dolejska M, Literak I, Fuchs S, Ahmed N, Grobbel M, Torres C, McNally A, Pickard D, Ewers C, Croucher NJ, Corander J, Guenther S. 2019. Genomic and Functional Analysis of Emerging Virulent and Multidrug-Resistant *Escherichia coli* Lineage Sequence Type 648. Antimicrob Agents Chemother 63:e00243–19.

15. McGowan JE, Barnes MW, Finland M. 1975. Bacteremia at Boston City Hospital: occurrence and mortality during 12 selected years (1935-1972), with special reference to hospital acquired cases. J Infect Dis 132:316–335.

16. Gransden WR, Eykyn SJ, Phillips I, Rowe B. 1990. Bacteremia Due to *Escherichia coli*: A Study of 861 Episodes. Rev Infect Dis 12:1008–1018.

17. Kallonen T, Brodrick HJ, Harris SR, Corander J, Brown NM, Martin V, Peacock SJ, Parkhill J. 2017. Systematic longitudinal survey of invasive *Escherichia coli* in England demonstrates a stable population structure only transiently disturbed by the emergence of ST131. Genome Res 27:1437–1449.

18. Laupland KB. 2013. Incidence of bloodstream infection: a review of population-based studies. Clin Microb Infect 19:492–500.

19. Laupland KB, Church DL. 2014. Population-based epidemiology and microbiology of community-onset bloodstream infections. Clin Microb Rev 27:647–664.

20. Olesen B, Scheutz F, Menard M, Skov MN, Kolmos HJ, Kuskowski MA, Johnson JR. 2009. Three-decade epidemiological analysis of *Escherichia coli* O15:K52:H1. J Clin Microb 47:1857–1862.

21. Rodríguez_Baño J, Picón E, Gijón P, Hernández JR, Ruíz M, Peña C, Almela M, Almirante B, Grill F, Colomina J, Giménez M, Oliver A, Horcajada JP, Navarro G, Coloma A, Pascual A. 2010. Community_Onset Bacteremia Due to Extended_Spectrum βLactamase–Producing *Escherichia coli*: Risk Factors and Prognosis. Clin Infect Dis 50:40–48.

22. CDC. Antibiotic Resistance Threats in the United States, 2013. Atlanta, GA: U.S. Department of Health and Human Services, CDC; 2013.

23. Merino I, Hernández-García M, Turrientes M-C, Pérez-Viso B, Ló Pez-Fresne∼ Na N, Diaz-Agero C, Maechler F, Fankhauser-Rodriguez C, Kola A, Schrenzel J, Harbarth S, Bonten M, Gastmeier P, Canton R, Ruiz-Garbajosa P. Emergence of ESBL-producing *Escherichia coli* ST131-C1-M27 clade colonizing patients in Europe. J Antimicrob Chemother 73:2973–2980.

24. Nicolas-Chanoine MH, Bertrand X, Madec JY. 2014. *Escherichia coli* ST131, an intriguing clonal group. Clin Microb Rev 27:543–574.

25. Rodríguez I, Novais Â, Lira F, Valverde A, Curião T, Martínez JL, Baquero F, Cantón R, Coque TM. 2015. Antibiotic-resistant *Klebsiella pneumoniae* and *Escherichia coli* high-risk clones and an IncFII(k) mosaic plasmid hosting Tn*1* (blaTEM-4) in isolates from 1990 to 2004. Antimicrob Agents Chemother 59:2904–8.

26. Novais A, Baquero F, Machado E, Cantón R, Peixe L, Coque TM. 2010. International spread and persistence of TEM-24 is caused by the confluence of highly penetrating enterobacteriaceae clones and an IncA/C2 plasmid containing Tn*1696*::Tn*1* and IS*5075*-Tn*21*. Antimicrob Agents Chemother 54:825–34.

27. Kallonen T, Brodrick HJ, Harris SR, Corander J, Brown NM, Martin V, Peacock SJ, Parkhill J. 2017. Systematic longitudinal survey of invasive *Escherichia coli* in England demonstrates a stable population structure only transiently disturbed by the emergence of ST131.Genome Res 27:1437–49.

28. Faith JJ, Colombel J, Gordon JI. 2014. Identifying strains that contribute to complex diseases through the study of microbial inheritance. Proc Natl Acad Sci U S A. 112:633–40.

29. Yamaji R, Rubin J, Thys E, Friedman CR, Riley LW. 2018. Persistent pandemic lineages of uropathogenic *Escherichia coli* in a college community from 1999 to 2017. J Clin Microbiol 56:e01834–17.

30. Olesen SW, Lipsitch M, Grad YH. 2019. The potential for “spillover” in outpatient antibiotic stewardship interventions among US states. bioRxiv 536714.

31. Low M, Neuberger A, Hooton TM, Green MS, Raz R, Balicer RD, Almog R. 2019. Association between urinary community-acquired fluoroquinolone-resistant *Escherichia coli* and neighbourhood antibiotic consumption: a population-based case-control study. Lancet Infect Dis 19:419–428.

32. Gottesman B-S, Low M, Almog R, Chowers M. 2019. Quinolone Consumption by Mothers Increases Their Children’s Risk of Acquiring Quinolone-Resistant Bacteriuria. Clin Infect Dis 71:532–538.

33. Fuzi M, Szabo D, Csercsik R. 2017. Double-serine fluoroquinolone resistance mutations advance major international clones and lineages of various multi-drug resistant bacteria. Front Microbiol 8:2261.

34. Redgrave LS, Sutton SB, Webber MA, Piddock LJV. 2014. Fluoroquinolone resistance: Mechanisms, impact on bacteria, and role in evolutionary success. Trends Microbiol 22:438–45

35. Yelin I, Snitser O, Novich G, Katz R, Tal O, Parizade M, Chodick G, Koren G, Shalev V, Kishony R. 2019. Personal clinical history predicts antibiotic resistance of urinary tract infections. Nat Med 25:1143–1152.

36. Price LB, Hungate BA, Koch BJ, Davis GS, Liu CM. 2017. Colonizing opportunistic pathogens (COPs): The beasts in all of us. PLoS Pathog 13:e1006369.

37. Kidsley AK, O’Dea M, Saputra S, Jordan D, Johnson JR, Gordon DM, Turni C, Djordjevic SP, Abraham S, Trott DJ. 2020. Genomic analysis of phylogenetic group B2 extraintestinal pathogenic *E. coli* causing infections in dogs in Australia. Vet Microbiol 248:108783.

38. Kidsley AK, O’Dea M, Ebrahimie E, Mohammadi-Dehcheshmeh M, Saputra S, Jordan D, Johnson JR, Gordon D, Turni C, Djordjevic SP, Abraham S, Trott DJ. 2020. Genomic analysis of fluoroquinolone-susceptible phylogenetic group B2 extraintestinal pathogenic Escherichia coli causing infections in cats. Vet Microbiol 245:108685.

39. Liu CM, Stegger M, Aziz M, Johnson TJ, Waits K, Nordstrom L, Gauld L, Weaver B, Rolland D, Statham S, Horwinski J, Sariya S, Davis GS, Sokurenko E, Keim P, Johnson JR, Price LB. 2018. *Escherichia coli* ST131-H22 as a foodborne uropathogen. mBio 9:e00470–18

40. Baquero F, Coque TM. 2011. Multilevel population genetics in antibiotic resistance. FEMS Microbiol Rev 35:705–706.

41. Selander RK, Levin BR. 1980. Genetic diversity and structure in *Escherichia coli* populations. Science 210:545–547.

42. Holt RD. 2009. Bringing the Hutchinsonian niche into the 21st century: Ecological and evolutionary perspectives. Proc Natl Acad Sci U S A. 106:19659–19665.

43. Thänert R, Reske KA, Hink T, Wallace MA, Wang B, Schwartz DJ, Seiler S, Cass C, Burnham CAD, Dubberke ER, Kwon JH, Dantas G. 2019. Comparative genomics of antibiotic-resistant uropathogens implicates three routes for recurrence of urinary tract infections. mBio 10.:e01977–19.

44. Bolnick DI, Svanbäck R, Araújo MS, Persson L. 2007. Comparative support for the niche variation hypothesis that more generalized populations also are more heterogeneous. Proc Natl Acad Sci U S A. 104:10075–10079.

45. Baquero F, Coque TM, Galán JC, Martinez JL. 2021. The Origin of Niches and Species in the Bacterial World. Front Microbiol 12:657986.

46. Sarkar S, Hutton ML, Vagenas D, Ruter R, Schüller S, Lyras D, Schembri MA, Totsika M. 2018. Intestinal Colonization Traits of Pandemic Multidrug-Resistant *Escherichia coli* ST131. J Infect Dis 218:979–990.

47. McNally A, Kallonen T, Connor C, Abudahab K, Aanensen DM, Horner C, Peacock SJ, Parkhill J, Croucher NJ, Corander J. 2019. Diversification of colonization factors in a multidrug-resistant *Escherichia coli* lineage evolving under negative frequency-dependent selection. mBio 10:e00644–19.

48. Amarsy R, Guéret D, Benmansour H, Flicoteaux R, Berçot B, Meunier F, Mougari F, Jacquier H, Pean de Ponfilly G, Clermont O, Denamur E, Teixeira A, Cambau E. 2019. Determination of *Escherichia coli* phylogroups in elderly patients with urinary tract infection or asymptomatic bacteriuria. Clin Microb Infect 25:839–844.

49. Anderson RM. 1999. The pandemic of antibiotic resistance. Nat Med 5:147–9.

50. Horcajada JP, Shaw E, Padilla B, Pintado V, Calbo E, Benito N, Gamallo R, Gozalo M, Rodr Iguez-Ba∼ No J. 2013. Healthcare-associated, community-acquired and hospital-acquired bacteraemic urinary tract infections in hospitalized patients: a prospective multicentre cohort study in the era of antimicrobial resistance. Clin Microb Infect 19:962–968.

51. National Committee for Clinical Laboratory Standards (CLSI). 2013. Performance Standards for Antimicrobial susceptibility Testing; Twenty-Third Informational Supplement. NCCLS, Wayne, PA, USA.

52. Clermont O, Christenson JK, Denamur E, Gordon DM. 2013. The Clermont *Escherichia coli* phylo-typing method revisited: Improvement of specificity and detection of new phylo-groups. Environ Microbiol Rep 5:58–65.

53. Clermont O, Christenson JK, Daubié AS, Gordon DM, Denamur E. 2014. Development of an allele-specific PCR for *Escherichia coli* B2 sub-typing, a rapid and easy to perform substitute of multilocus sequence typing. J Microbiol Methods 101:24–27.

54. Johnson JR, Clermont O, Johnston B, Clabots C, Tchesnokova V, Sokurenko E, Junka AF, Maczynska B, Denamur E. 2014. Rapid and specific detection, molecular epidemiology, and experimental virulence of the O16 subgroup within Escherichia coli sequence type 131. J Clin Microbiol 52:1358–1365.

55. Russo TA, Johnson JR. 2000. Proposal for a New Inclusive Designation for Extraintestinal Pathogenic Isolates of *Escherichia coli*: ExPEC. J Infect Dis 181:1753–1754.

56. Bankevich A, Nurk S, Antipov D, Gurevich AA, Dvorkin M, Kulikov AS, Lesin VM, Nikolenko SI, Pham S, Prjibelski AD, Pyshkin A v., Sirotkin A v., Vyahhi N, Tesler G, Alekseyev MA, Pevzner PA. 2012. SPAdes: A New Genome Assembly Algorithm and Its Applications to Single-Cell Sequencing. J Comput Biol 19:455.

57. Seemann T. 2014. Prokka: rapid prokaryotic genome annotation. Bioinformatics 30:2068–2069.

58. Fernández-de-Bobadilla MD, Talavera-Rodríguez A, Chacón L, Baquero F, Coque TM, Lanza VF. 2021. PATO: Pangenome Analysis Toolkit. Bioinformatics btab697.

59. Price MN, Dehal PS, Arkin AP. 2010. FastTree 2 – Approximately Maximum-Likelihood Trees for Large Alignments. PLoS One 5:e9490.

60. Yu G, Smith DK, Zhu H, Guan Y, Lam TT-Y. 2017. ggtree: an r package for visualization and annotation of phylogenetic trees with their covariates and other associated data. Meth Ecol Evol 8:28–36.

