## Supplementary material for "A 21-year survey of *Escherichia coli* from bloodstream infections (BSIs) in a tertiary hospital reveals how community-hospital dynamics of the B2 phylogroup clones influence local BSI rates": 20211102_Supplementary Text_BSIcoli_SENT_mSphere 2021 copy.docx

**Virulence-associated gene profiling in the B2 phylogroup of *Escherichia coli***

The presence of 29 virulence-associated genes (VAGs) was determined for all B2 isolates by PCR (1, 2). The B2 *E. coli* strains were clustered into 2 large groups and 10 subgroups on the basis of the content of the VAG genes. Each group comprises numerous gene combinations and showed variable redundancies. Most *E. coli* isolates analyzed here harbored *fyuA,* an outer membrane protein acting as a receptor for ferric-yersiniabactin (*fyu*) and a marker of PAIJ86IV (a “high pathogenicity island”) (3, 4); *malX,* a phosphotransferase system enzyme and marker of PAICFT073 (5); *ompT* (an extracellular protease active against host cationic peptides located on *E. coli* plasmids or chromosomes (6, 7); and *traT*, only located on F plasmids, which act as a surface exclusion protein, conferring serum resistance. (8). However, the isolates differ in the presence of type P fimbriae, all of which contain various combinations of siderophores, toxins, and types S and I fimbriae, which are usually located on emblematic *E. coli* pathogenicity islands (PAIs). Despite some VAG variability within each B2 subgroup, more than half (56%) of the B2-I (STc131) strains clustered in VAG 1.5. The isolates of the *E. coli* phylogenetic subgroups B2_II_, B2_III,_ B2_V_, B2_VII_, B2_IX_, and B2_X_ clustered in the VAG 2.6, 2.2, 2.9, 2.6, 2.7, and 2.4 groups, respectively (54%–75%) (Figures 5, S4, S5, S6, S7). The consistent association between B2 subgroups and VAG profiles may be influenced by the presence of clonally related isolates within each subgroup (e.g. B2_II_, B2_IX,_ B2_I_-ST131).

The VAG 1.5 group only comprised isolates of the B2-I (STc131) and B2-VII subgroups (representing 54% and 33% of the isolates of these B2 subgroups, respectively). They contained the genes *iha, sat, iutA,* *kpsMT,* and sporadically *hlyA*, all frequently located on PAICFT073-pheV*,* and to a lesser extent, *afa/draBC* (25%) or *ibeA*. Some B2-I (STc131) isolates were also associated with VAG 1.8 (lacking traits of PAICFT073-pheV but containing *ibeA* and often *iroN*), VAG 2.4 (containing *iroN*, *ibeA,* and *papABCED*), and VAG2.9 (*papABCED)*. The analysis of STc131 isolates in Figure S7 reflects the diversity of VAGs within the ST131 groups H30 (or subclade C), H22 (or subclade B), and H41 (or subclade A); many of the isolates exhibiting VAGs could not be associated with any of the virotypes proposed by Dahbi et al. (9)

The VAG 2.7 was predominant in isolates of the B2-IX group (63%, p <.005) and was only associated with this B2 subgroup. This VAG profile was characterized by *papACEF* genes, *ireA* (encoding a protein involved in iron acquisition, adhesion to epithelial cells, resistance to stress, and a high pH, a trait frequently associated with isolates causing UTIs and BSIs), and a variable presence of genes in PAIU189 (*iss*, *cvaC*, and K1-capsule).

The VAG 2.6 profile comprised more than half of B2-II and B2-VII isolates and was linked to the presence of type P fimbriae and with various traits associated with PAIJ96 (*hlyA, cnf1, sfa, ΔpapGIII*), PAICFT073-serX *(sfa/focDE, iroN*), and PAICFT073-pheV (*iutA, iha, iroN, sat, kpsMTK5*).

The VAG 2.9 group was mostly associated with B2-V (50% of the isolates), and differs from VAG 2.6 in the absence of *iroN* and *K5*. VAG 2.2 comprises most B2-III (73 of the isolates) and VAG 2.4, most B2-X isolates. With the exception of B2-I and B2-IV, genes of most strains of B2-II, B2-IX, and B2-IV have traditionally been associated with UTIs or BSIs.

**References (Supplementary Text)**

1.	Johnson JR, Russo TA. 2018. Molecular Epidemiology of Extraintestinal Pathogenic *Escherichia coli*. EcoSal Plus 8.

2.	Russo TA, Johnson JR. 2000. Proposal for a New Inclusive Designation for Extraintestinal Pathogenic Isolates of *Escherichia coli*: ExPEC . *J Infect Dis* 181:1753–1754.

3.	Clermont O, Bonacorsi S, Bingen E. 2001. The Yersinia high-pathogenicity island is highly predominant in virulence-associated phylogenetic groups of *Escherichia coli* . *FEMS Microbiol Letters* 196:153–157.

4.	Hancock V, Ferrières L, Klemm P. 2008. The ferric yersiniabactin uptake receptor FyuA is required for efficient biofilm formation by urinary tract infectious *Escherichia coli* in human urine. *Microbiology* 154:167–175.

5.	Östblom A, Adlerberth I, Wold AE, Nowrouzian FL. 2011. Pathogenicity island markers, virulence determinants malX and usp, and the capacity of *Escherichia coli* to persist in infants’ commensal microbiotas. *Applied Environm Microbiol* 77:2303–2308.

6.	Stumpe S, Schmid R, Stephens DL, Georgiou G, Bakker EP. 1998. Identification of OmpT as the protease that hydrolyzes the antimicrobial peptide protamine before it enters growing cells of *Escherichia coli*. *J. Bacteriol* 180:4002–4006.

7.	Dan M, Yair Y, Samosav A, Gottesman T, Yossepowitch O, Harari-Schwartz O, Tsivian A, Schreiber R, Gophna U. 2015. *Escherichia coli* isolates from patients with bacteremic urinary tract infection are genetically distinct from those derived from sepsis following prostate transrectal biopsy. *Int J Med Microbiol* 305:464–468.

8.	Harrison JL, Taylor IM, Platt K, O’Connor CD. 1992. Surface exclusion specificity of the TraT lipoprotein is determined by single alterations in a five‐amino‐acid region of the protein. *Mol Microbiol* 6:2825–2832.

9.	Dahbi G, Mora A, Mamani R, López C, Alonso MP, Marzoa J, Blanco M, Herrera A, Viso S, García-Garrote F, Tchesnokova V, Billig M, de la Cruz F, de Toro M, González-López JJ, Prats G, Chaves F, Martínez-Martínez L, López-Cerezo L, Denamur E, Blanco J. 2014. Molecular epidemiology and virulence of *Escherichia coli* O16: H5-ST131: Comparison with H30 and H30-Rx subclones of O25b: H4-ST131. *Int J Med Microbiol* 304:1247–1257.
