## Supplementary figures and images for "A 21-year survey of *Escherichia coli* from bloodstream infections (BSIs) in a tertiary hospital reveals how community-hospital dynamics of the B2 phylogroup clones influence local BSI rates"

### 20211103_Supplementary figures_BSIcoli_OK_SENT copy.pdf

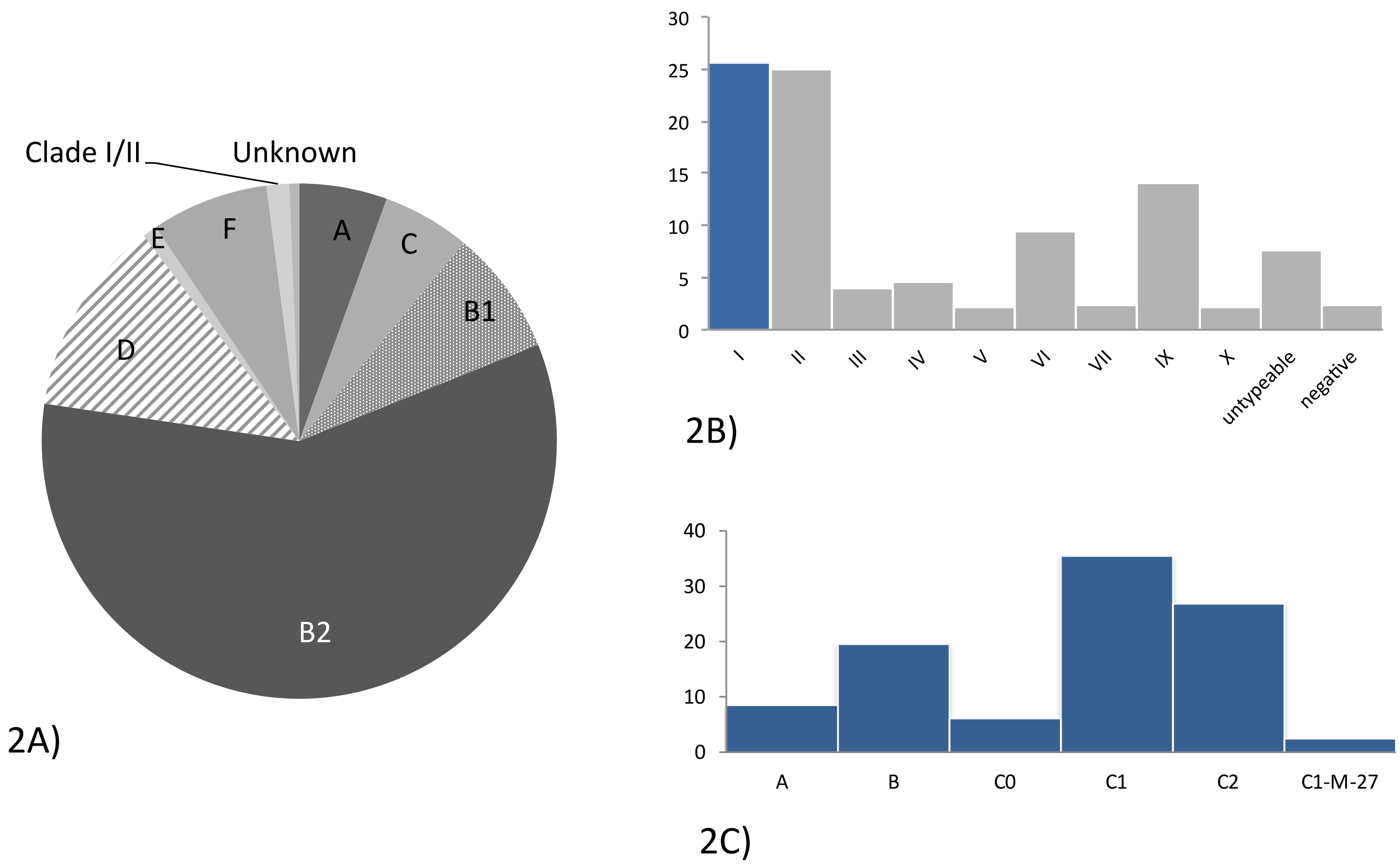

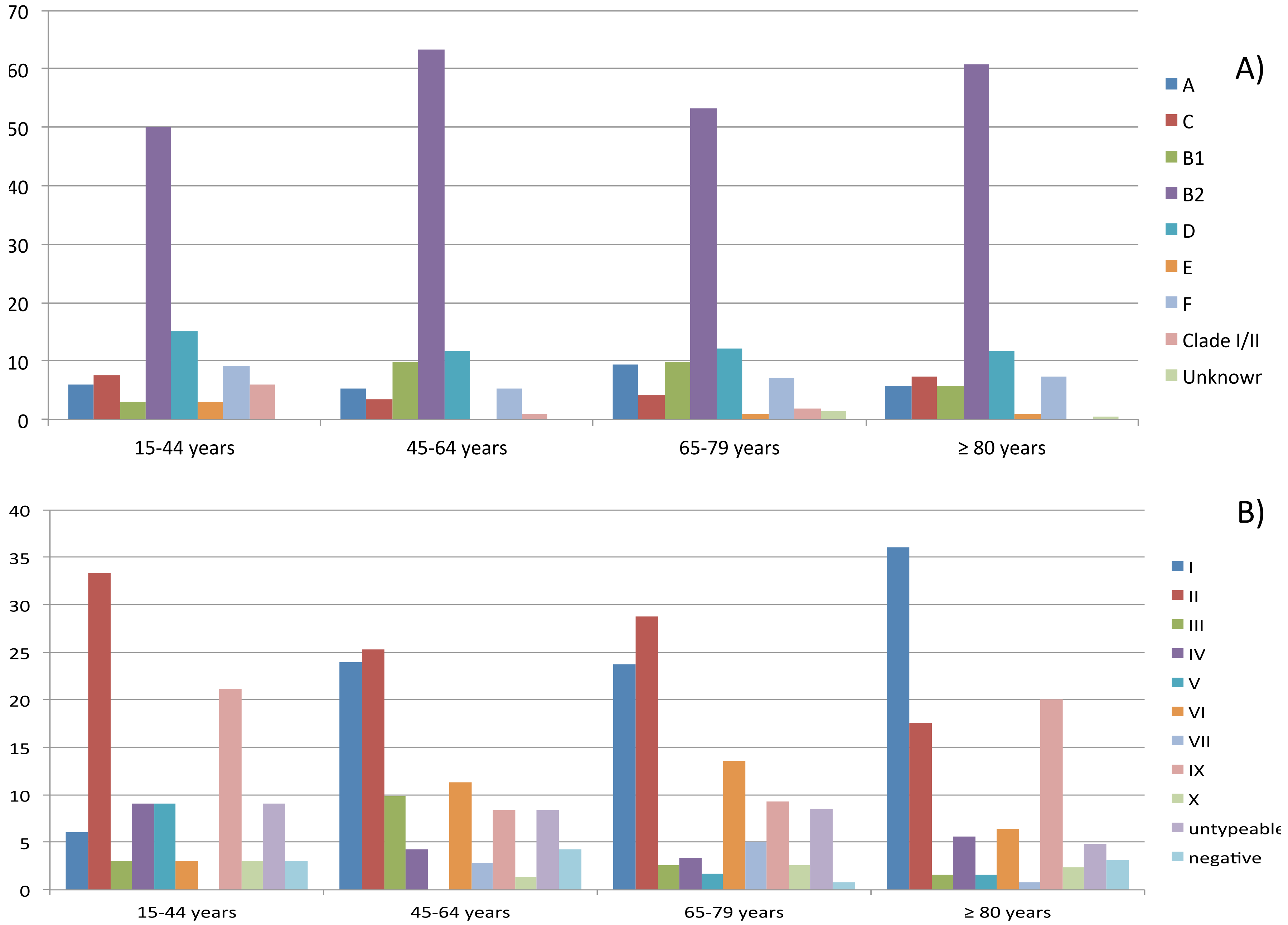

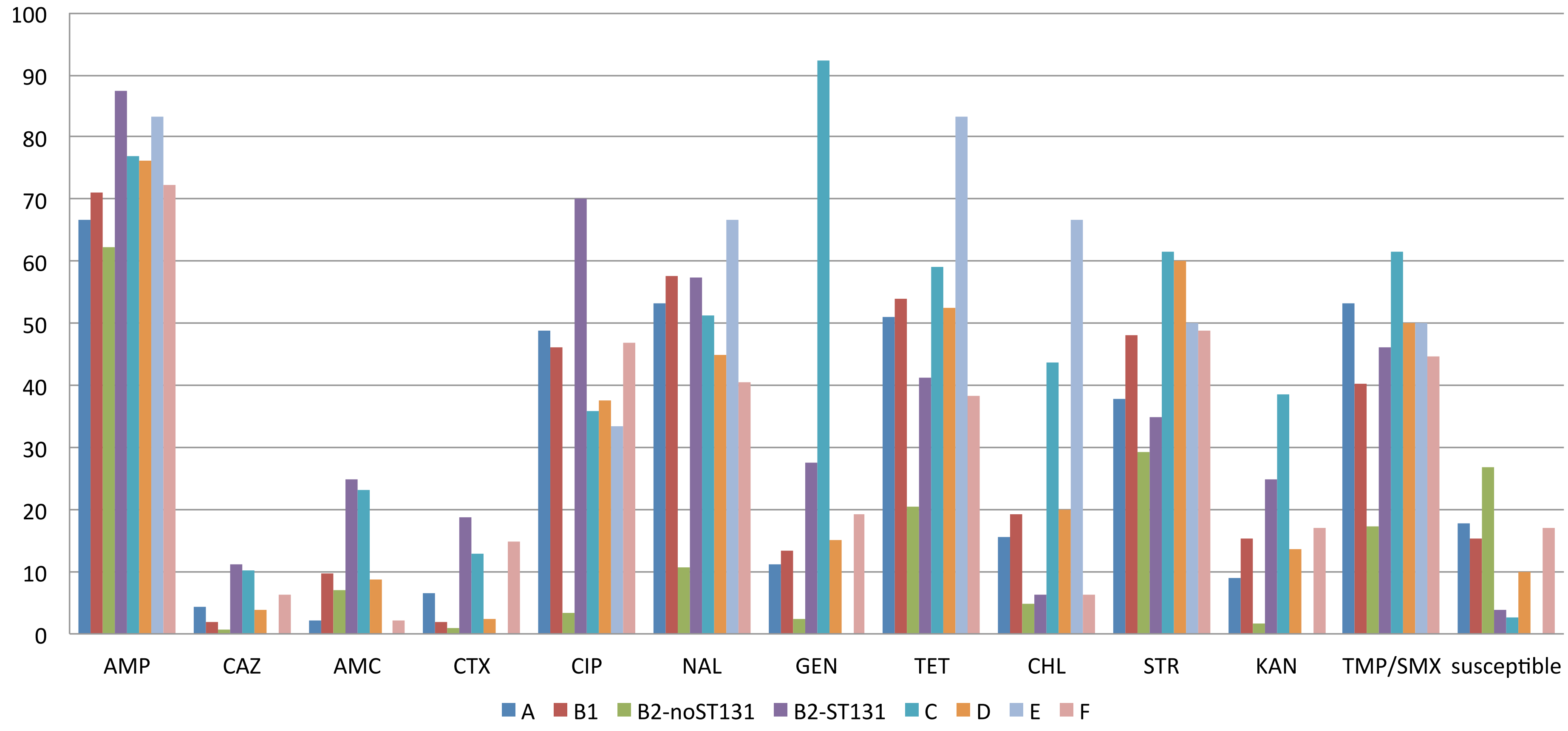

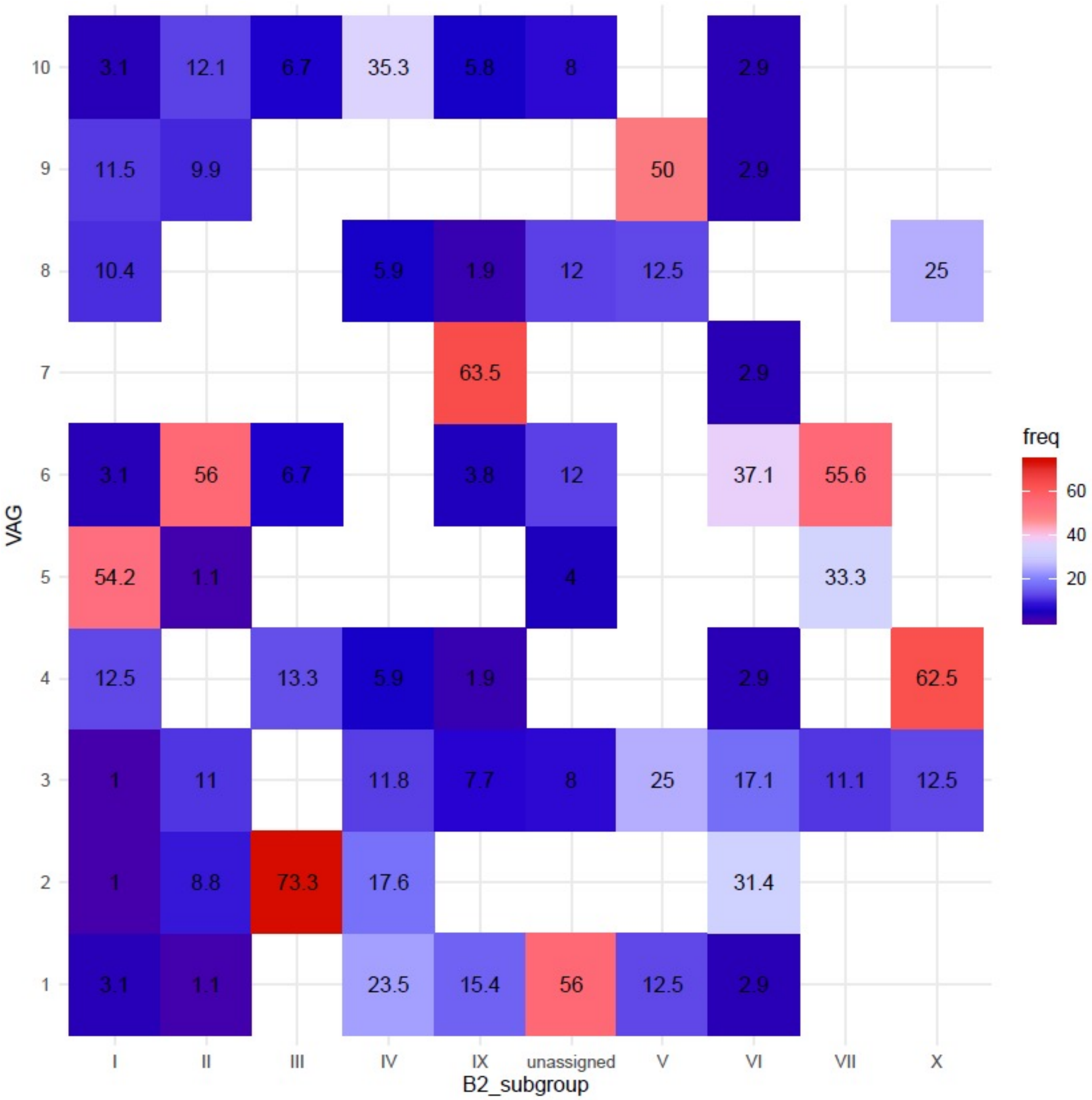

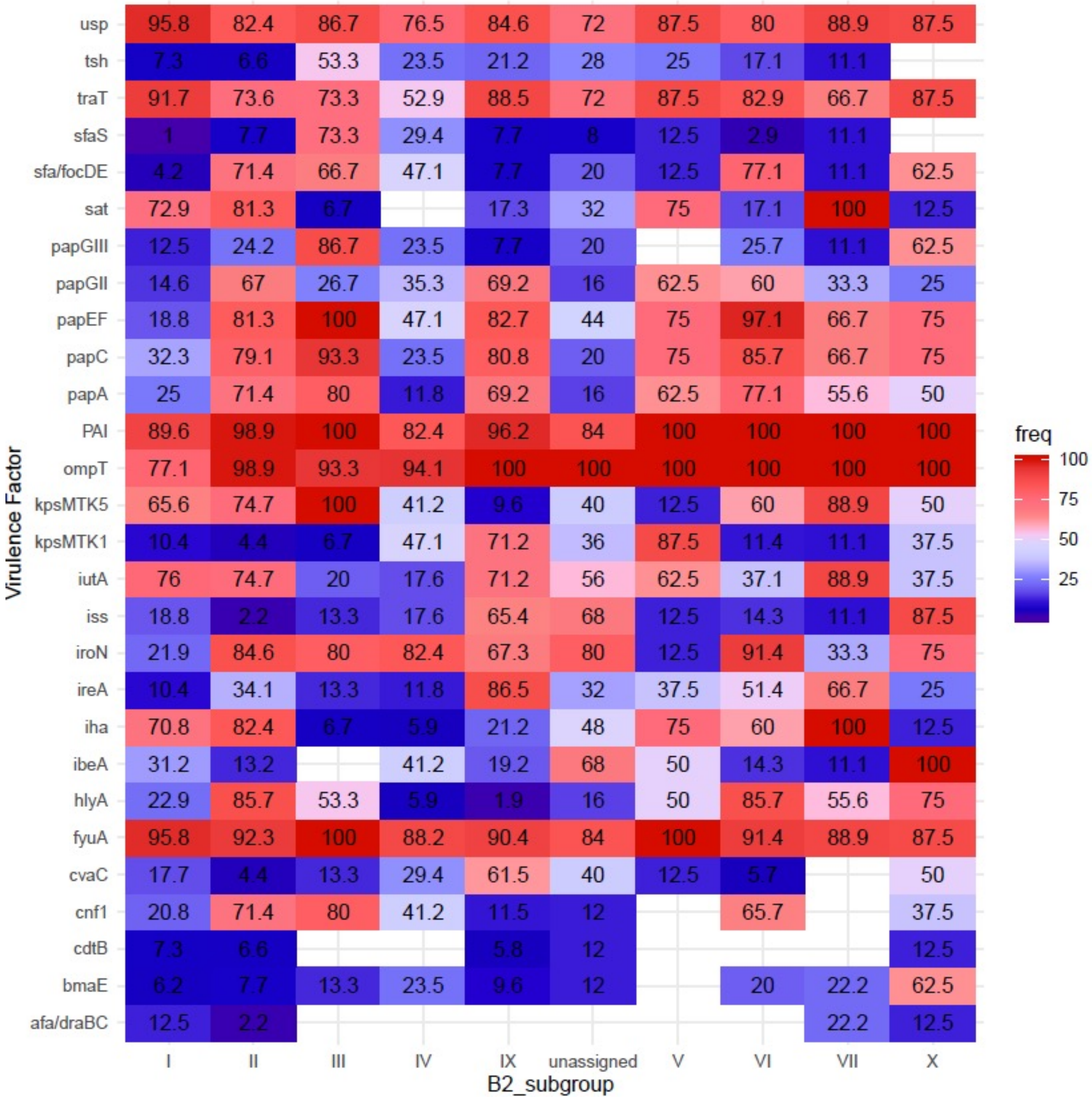

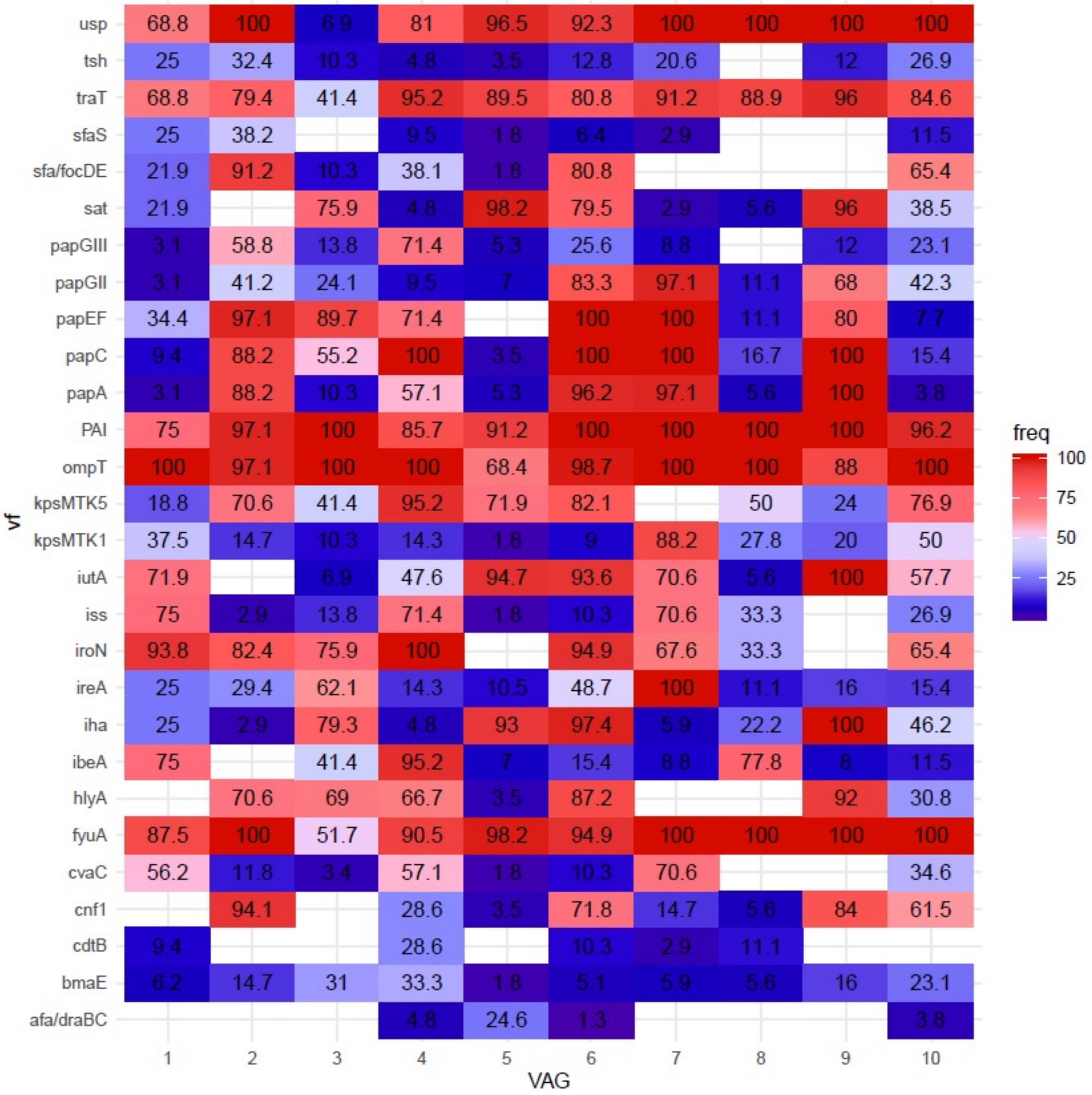

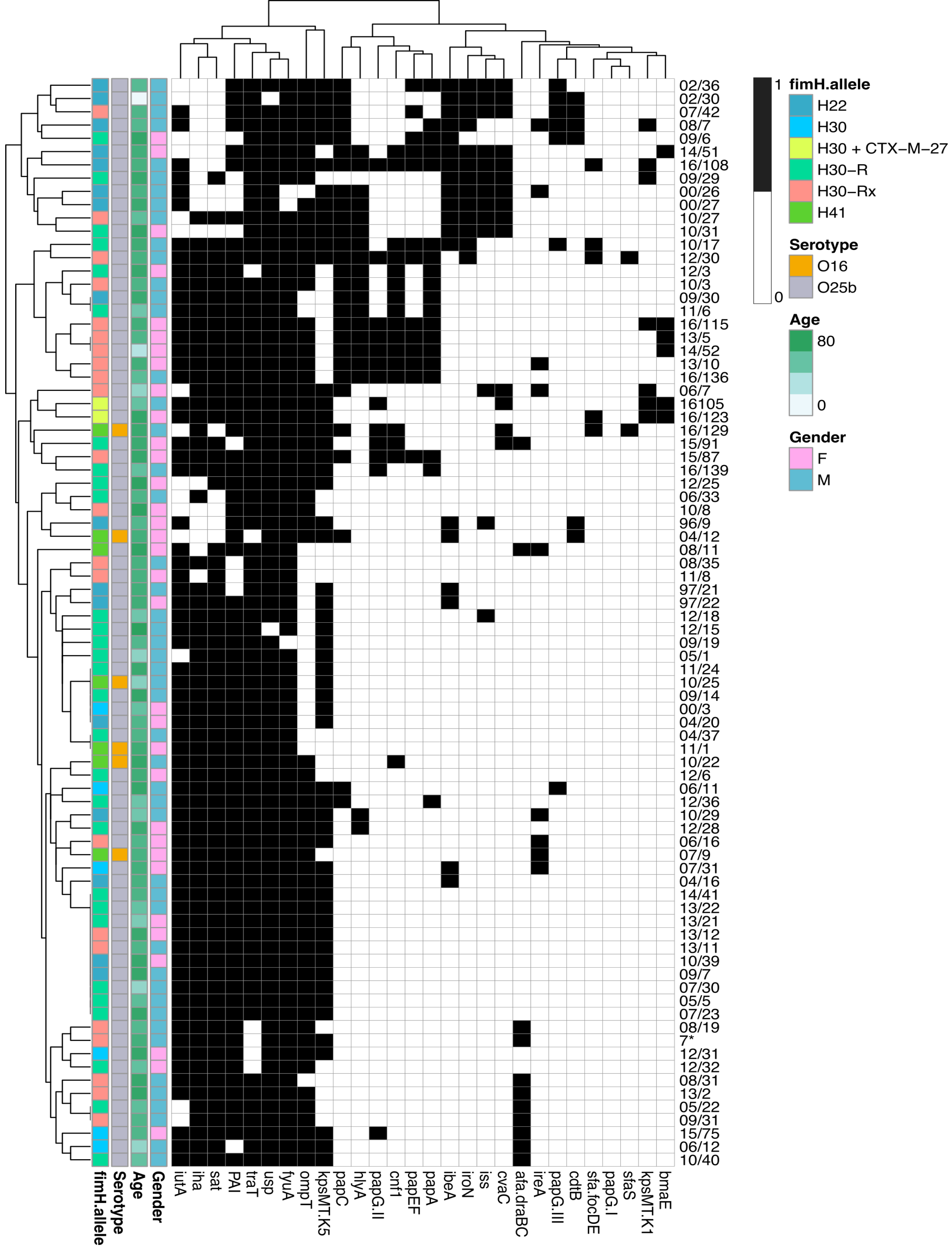
